# BARTsc identifies key transcriptional regulators from single-cell omics data

**DOI:** 10.64898/2026.02.24.707729

**Authors:** Hongpan Zhang, Lei Kang, Jingyi Wang, Kathleen P. Liang, Zhenjia Wang, Kexin Xu, Chongzhi Zang

**Affiliations:** Department of Genome Sciences, University of Virginia, Charlottesville, VA 22908, USA; Department of Biochemistry and Molecular Genetics, University of Virginia, Charlottesville, VA 22908, USA; Department of Biology, University of Virginia, Charlottesville, VA 22908, USA; UVA Comprehensive Cancer Center, University of Virginia, Charlottesville, VA 22908, USA; Department of Biomedical Engineering, University of Virginia, Charlottesville, VA 22908, USA

## Abstract

Inference of transcriptional regulatory mechanisms from single-cell (sc) omics data, such as scRNA-seq, scATAC-seq, and scMultiome, remains an important problem in single-cell biology and functional genomics. Most existing methods for predicting functional transcriptional regulators (TRs) from single-cell data rely on co-expression between regulator and target genes and/or sequence motif enrichment, holding inherent limitations. Here, we present BARTsc, a computational method that accurately predicts functional TRs from clustered single-cell omics data by leveraging a large collection of public ChIP-seq profiles. BARTsc implements a novel framework to infer a cis-regulatory profile from differential genomic features from either unimodal (RNA or ATAC) or bimodal (scMultiome) single-cell profiling data and identify TRs whose binding profiles most associate with the cis-regulatory profile. BARTsc can quantify TR activity across cell clusters and predict key regulators for each cell cluster. We demonstrate that BARTsc can successfully identify active TRs in each cell type and cell-type-defining key regulators across diverse biological systems, including mouse cortex, human peripheral blood mononuclear cells (PBMCs), and human pancreatic ductal adenocarcinoma (PDAC). Using a generative-AI-assisted, literature-supported collection of cell-type key regulators as benchmarks, we show that BARTsc consistently outperforms existing state-of-the-art methods. We apply BARTsc to identify critical regulators in PDAC, including NEFLA, a novel PDAC key regulator, and validate its function in pancreatic tumor proliferation by experiments. As a robust and versatile computational method, BARTsc provides deeper insights into cell-type-specific regulatory programs, facilitating the discovery of key regulators across diverse biological systems.

## Introduction

Transcriptional regulators (TRs), including sequence-specific transcription factors (TFs) and chromatin regulators, play crucial roles in transcriptional regulation and are implicated in numerous cellular processes, including cancer development and progression^1–3^. The differential activity of TRs can lead to destiny-shaping changes in cell states. For example, upregulation of a group of TRs, such as ZEB1/2, SNAI1, SNAI2 and TWIST induces epithelial-mesenchymal transition (EMT), which enables tumor metastasis^4,5^. However, identifying which TRs are active within a cell system is challenging, as they can remain functionally effective even at low expression levels, making gene expression profiling alone insufficient to determine their activity. Meanwhile, existing experimental methods that assay TR binding activity (e.g., ChIP-seq^6^, CUT&RUN^7^, CUT&Tag^8^, etc.) are limited to examining only one factor at a time, making it unfeasible to comprehensively assess a large number of TRs within a given cell system. Therefore, inferring functional TRs from available data, such as transcriptomic profiles and/or chromatin accessibility profiles, has become an important task for studying gene regulation.

Recently developed single-cell genomics techniques^9–12^, including single-cell RNA-seq (scRNA-seq) and single-cell ATAC-seq (scATAC-seq), offer key advantages for studying transcriptional regulation: First, they capture cellular heterogeneity rather than a bulk average across multiple distinct cell states. Second, they clearly identify distinct cell types with different transcriptomic profiles, which are likely regulated by different TRs. Third, beyond scRNA-seq, scATAC-seq provides chromatin accessibility information on the single cell level, and scMultiome (ATAC+RNA) can profile gene expression and chromatin accessibility in the same cell. These more advanced techniques allow for the joint use of both transcriptomic and epigenomic data to infer regulatory mechanisms, and thus underscore the need for computational methods capable of identifying functional TRs from single-cell genomics and multiomics data.

All the existing computational methods for transcriptional regulation inference from single-cell omics data face the same challenges of establishing regulatory relationships between TRs and their target genes. Conventional approaches address this either by assessing TF binding motif (TFBM) enrichment in gene-proximal regions^13,14^, or by evaluating marginal associations such as TF-target co-expression^15,16^, while some employ hybrid strategies that leverage both approaches^17–19^. These approaches have inherent limitations: (1) TFBMs do not represent true binding sites —TFBM-based methods may yield both false positives due to >90% of unbound motifs^20^ and false negatives from frequent motif-less binding^21,22^; (2) focusing on gene-proximal regions overlooks distal enhancers, which play significant roles in transcriptional regulation in mammalian cells^23^; (3) TF-target co-expression can be masked by confounding from other TFs and shared regulatory programs^24^, and it only indicates correlation rather than causation. Moreover, many TRs exert strong regulatory effects even at low expression levels. New approaches are being explored to address these concerns. Some methods begin with prior binding knowledge and learn the weights of regulatory relationships by optimizing a Bayesian framework^25,26^, while some are based on in-silico perturbation^27,28^.

To address these challenges, we previously developed BART, a method for predicting functional TRs from a gene set using published TR ChIP-seq profiles as reference data^29–31^. BART uses a model-based approach to infer a genome-wide cis-regulatory profile from an input feature list (e.g., differentially expressed genes (DEGs) between two conditions usually derived from bulk RNA-seq) and estimates TR binding activity by quantifying the association between the genome-wide cis-regulatory profile and ChIP-seq profiles. The performance of BART has been validated through multiple studies and independent benchmarking works^32–35^. However, although BART has been utilized by the community in single-cell data analysis^36^, BART was originally designed for bulk sequencing data and does not account for the presence of multiple distinct cell types in single-cell sequencing data. Meanwhile, the emergence of single-cell multiome profiling technologies which simultaneously assay transcriptomics and chromatin accessibility presents new opportunities for a better understanding of transcriptional regulatory events driven by TR binding at chromatin. Therefore, built upon our existing BART framework, we sought to develop a new method that better leverages the heterogeneity information in single-cell omics data to infer transcriptional regulation and to integrate multiomic information for higher accuracy and biological resolution.

Here, we introduce BARTsc, a new method for inference of functional TRs that translate into differential cell states measured by single-cell omics assays. BARTsc can be applied to various data types as input, including scRNA-seq, scATAC-seq and single-cell Multiome (ATAC+RNA) data. We first describe the overall design of BARTsc, which consists of three distinct types of analyses, followed by a detailed evaluation of its performance using several real single-cell datasets across different scenarios. By comparing with several existing methods for identifying key regulators from different data types and different biological systems, we demonstrated that BARTsc is the best-performing method for identifying active TRs from single-cell omics data. Finally, we applied BARTsc to a single-cell Multiome dataset for human pancreatic ductal adenocarcinoma (PDAC) to identify novel TRs potentially driving pancreatic cancer development. We further validated the predictions experimentally, to confirm BARTsc’s ability to uncover novel TRs from single-cell data with significant biological relevance.

## Results

### BARTsc leverages differentially expressed features and public ChIP-seq data to predict TR binding activity

BARTsc is designed to take in data derived from multiple single-cell omics technologies, including scRNA-seq, scATAC-seq, and single-cell Multiome that simultaneously profile transcriptome and chromatin accessibility. Considering that a heterogeneous cell population from a single-cell dataset only contains a limited number of cell types and that single-cell sequencing data from individual cells is usually too sparse to capture sufficient biological information to distinguish one from another within a cell type or cell cluster, while imputation frequently introduce extra artifacts^37^, we design BARTsc to analyze single-cell omics data at the cell cluster or cell group level, where cells are grouped in clusters based on overall similarity or prior knowledge such as known cell type annotations (Fig. 1a).

**Figure 1.**
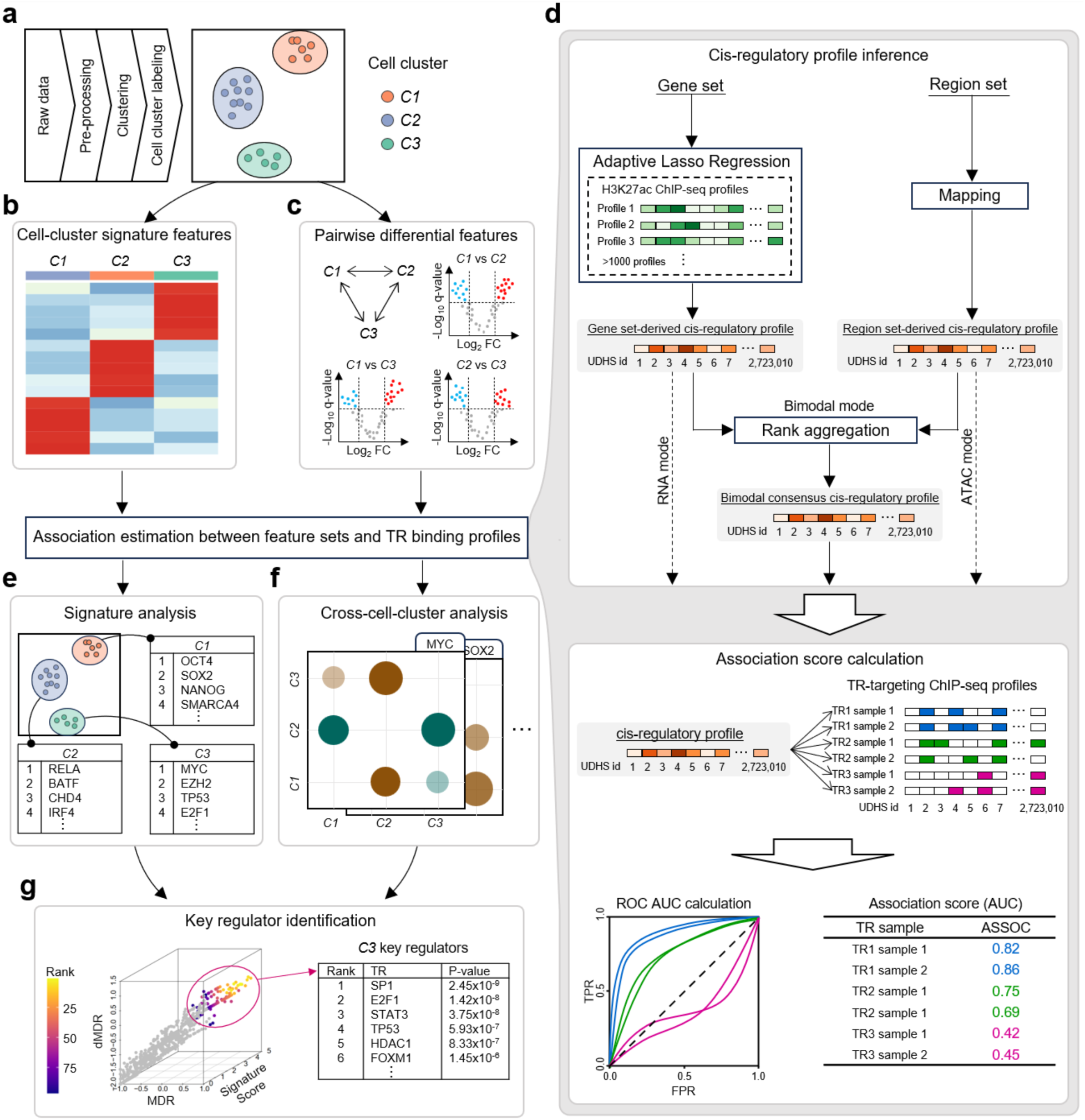
BARTsc Architecture. BARTsc takes a clustered single-cell dataset as input (**a**), identifies cell-cluster signature features (**b**) and pairwise differential features (**c**). To estimate the association between feature sets and published TR binding profiles, BARTsc first converts each feature set into a cis-regulatory profile using a modality-specific method, and then quantifies the association with each TR profile (**d**). BARTsc performs two parallel analyses: signature analysis (**e**) and cross-cluster analysis (**f**). BARTsc then integrates the results to identify key transcriptional regulators (**g**).

BARTsc first performs a feature extraction step to identify biologically relevant feature sets, such as differentially expressed genes or differentially accessible chromatin regions. Two distinct types of feature sets are simultaneously identified: cell-cluster signature feature sets, which comprise features specifically expressed or upregulated in each cell cluster compared to all the other clusters (Fig. 1b), and pairwise differential feature sets, which include features that are differential between every two distinct cell clusters (Fig. 1c).

To investigate the relationship between the biologically relevant feature sets and TR binding, BARTsc then quantifies the association between each identified feature set and known TR binding profiles (Fig. 1d). This process first infers a cis-regulatory profile for each feature set and then estimates the level of association between the inferred cis-regulatory profiles and known TR binding profiles^29,38^. Specifically, BARTsc uses union DNaseI hypersensitive sites (UDHS) as the basis for all potential cis-regulatory elements in the genome^38^. For feature sets derived from scRNA-seq, which are gene sets, BARTsc applies adaptive lasso regression to select and weight a subset from over 1,000 publicly available H3K27ac ChIP-seq profiles that best explains the expression pattern of the input gene set. The resulting weighted profile is defined as the cis-regulatory profile corresponding to the gene set. For feature sets derived from scATAC-seq, which are region sets, BARTsc directly maps the normalized peak signal to UDHSs to generate a cis-regulatory profile scored by chromatin accessibility level. For bimodal data such as scMultiome, a matched pair of gene set and chromatin region set is generated for each comparison. BARTsc infers a cis-regulatory profile from each modality separately and then uses a rank aggregation-based method to generate a bimodal consensus cis-regulatory profile, which highlights the regulatory elements supported by both transcriptomic and epigenomic signals. For efficient association estimation, all TR binding profiles are mapped to UDHS so that they can directly compare with cis-regulatory profiles. An association score is then calculated between each cis-regulatory profile and each TR binding profile as the area under the receiver operator characteristics curve (AUROC), which quantifies the predictive power of using the given cis-regulatory profile to classify the bound and unbound sites of the given TR.

The association scores derived from cell-cluster signature feature sets are used for a cell-cluster-centered signature analysis, which identifies the TRs that are most likely to explain the given cell cluster’s signature features (Fig. 1e). The association scores derived from pairwise differential feature sets are used for a TR-centered cross-cell-cluster analysis, which evaluates TRs’ relative activity across cell clusters by doing paired statistical tests between the association scores derived from the up-regulated feature set and those derived from the down-regulated feature set for each pair of cell clusters (Fig. 1f). Finally, by comprehensively integrating the TRs’ contribution to signature feature expression and their relative activity patterns within the given dataset, BARTsc identifies key regulators for each cell type in an unbiased way (Fig. 1g).

### Signature analysis Identified functional TRs from single-cell transcriptomics data

We first sought to validate the performance of BARTsc on a unimodal dataset. We applied BARTsc signature analysis to the six major cell types of a mouse cortex scRNA-seq dataset from a published study^39^ (Fig. 2a), a used single-cell dataset for computational method development. We observed that the identified TRs with statistical significance are with well-characterized functions in corresponding cell types (Fig. 2b, Fig. S1a-f). For example, TRs essential for the differentiation and function of astrocytes, such as SOX9, TCF4 and PPARγ^40–42^, had high signature scores in the signature analysis result for astrocytes. Meanwhile, these astrocyte-specific TRs received lower scores in other cell types, especially neurons. For oligodendrocytes, SOX10, an essential TR that restricts the differentiation of oligodendrocytes precursor cells (OPCs) toward the oligodendrocyte lineage^43^, together with another key regulator DICER^44^ received high signature scores. For microglia, well-known macrophage markers including SPI1(PU.1), RELA, IRF8, IRF3 and STAT3 were identified as top signature TRs^45–49^. A series of regulators that are generally critical for neuronal development and cell identity maintenance, such as NEUROD2, MEF2A, MEF2C, NPAS4, E2F1 and FGFR1^50–55^, all received high signature scores in the two neuronal cell types. Consistently, interneuron-specific regulators (MECP2^56^, PRDM13^57^) showed high scores in interneurons, while pyramidal cell-specific regulators (AUTS2^58^, CREB1^59^) were assigned high scores in pyramidal cells. Endothelial cells and mural cells were grouped together due to their mutualistic and essential relationship in the formation of blood vessels. TRs that participate in angiogenesis and vascular function, such as CREB1, FLI1, ETS1, ETV2, KLF4, GATA4, SRF, SMAD1, FOXO1 were scored high for endothelial cells and mural cells^60,61^. These results suggest that BARTsc can predict cell-type functional TRs that are critical for cell differentiation and the maintenance of cell identity based on signature genes derived from scRNA-seq data.

**Figure 2.**
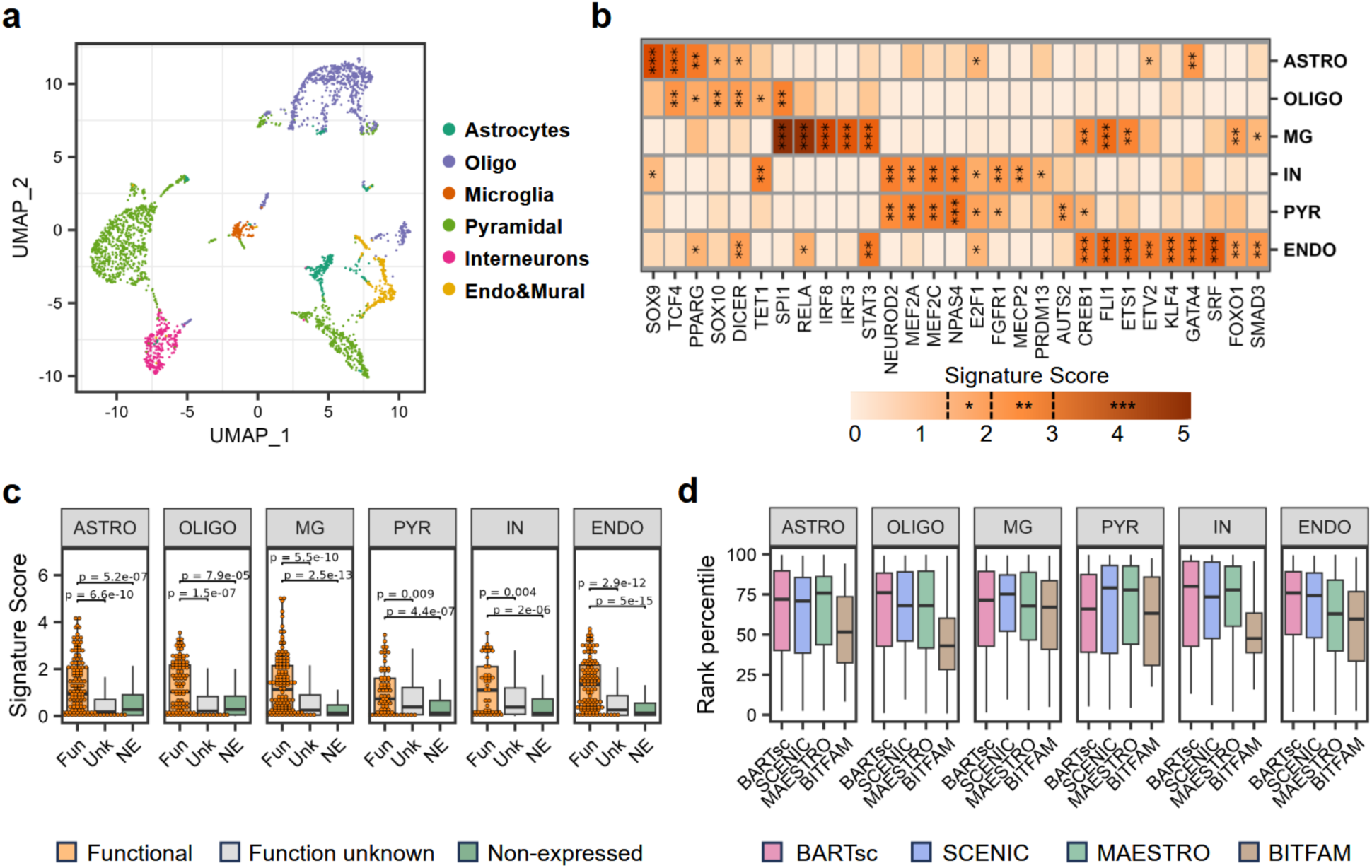
Signature analysis reveals functional TRs from scRNA-seq. (**a**) UMAP visualization of the scRNA-seq dataset for mouse cortex, where each point represents a single cell colored by cell type. (**b**) Signature analysis results for selected TRs in mouse cortex. The heat map is colored by signature score. Asterisks indicate significance level, *: *p* < 0.05, **: *p* < 0.01, ***: *p* < 0.001. (**c**) Box plots showing the distributions of signature scores of functional TRs (“Fun”, orange), function-unknown TRs (“Unk”, grey) and non-expressed TRs (“NE”, green) across different cell types. In each box plot, the center line represents median; the box denotes the interquartile range (IQR); the whiskers extend to the minimum and maximum values within 1.5× IQR from the median. All individual data points for functional TRs are overlaid on the boxes. P-values were calculated using one-sided Wilcoxon rank-sum tests. (**d**) Box plots showing the distributions of the rank percentile of functional TRs predicted by each method across different cell types. In each box, the center line represents median; the box denotes the interquartile range (IQR); the whiskers extend to the minimum and maximum values within 1.5× IQR from the median.

We next examined whether the signature scores generated by BARTsc can distinguish functional TRs from other TRs. To do this, we curated an extensive list of functional TRs for each cell type based on AI-assisted literature review (see Methods). We classified the TRs into three groups based on their RNA expression and function in each cell type: (1) the expressed TRs with known functions in the given cell type were labeled as functional TRs; (2) the expressed TRs without direct evidence of functionality in the given cell type were labeled as function-unknown TRs; and (3) the non-expressed TRs in a given cell type were labelled as non-expressed TRs. For each cell type, comparisons among the three TR groups indicated that the functional group has significantly higher signature scores than the other two groups (Fig. 2c), suggesting that the BARTsc signature scores effectively distinguish functional TRs from other TRs. We next benchmarked BARTsc against several published methods that output ranked TR lists based on scRNA-seq data. To ensure a fair comparison, we calculated the rank percentiles of functional TRs in each cell type for every method. Across the six cell types, BARTsc achieved the highest median rank percentile in three cell types and the second highest in two other cell types, representing the best overall performance among the benchmarked methods (Fig. 2d). Interestingly, BARTsc assigns lower rank percentiles to function-unknown TRs and the lowest rank percentiles to non-expressed TRs compared to other methods (Fig. S1g, h), suggesting that it achieves high sensitivity while maintaining high specificity. In conclusion, BARTsc exhibited high accuracy in identifying cell-type-functional TRs from scRNA-seq data and outperforms other existing methods.

### Integration of transcriptome and chromatin accessibility information improves TR prediction accuracy

With the application of the cutting-edge single-cell multiome technique, the transcriptome and chromatin accessibility profiles can be assayed simultaneously for the same individual cell. While transcription levels and chromatin accessibility are generally correlated, each provides specific biological information that cannot be fully represented by the other. Thus, we hypothesized that BARTsc can achieve superior prediction power by integrating both the transcriptome and chromatin accessibility information and prioritizing the regulatory regions informed by both modalities.

To test this hypothesis, we reanalyzed a 10X multiome (RNA+ATAC) dataset for healthy human PBMCs, a relatively well-studied system. Based on marker gene expression, we labeled 5 major cell types and 7 subtypes (Fig. 3a, Fig. S2). Applying BARTsc’s bimodal mode, we successfully identified known functional regulators associated for distinct immune cell lineages (Fig. 3b), including EBF1, PAX5, FOXP1, SPIB, and MEF2C in B cells^62–66^, MYC, RBPJ, NOTCH1, TCF7, STAT5A, and STAT5B in T cells^67–69^, TBX21, STAT4, and EOMES in NK cells^70,71^, CEBPA, CEBPB, SPI1 (PU.1), NR4A1, STAT1, and IRF1 in monocytes ^72–75^, and IRF4, IKZF1, SPI1 (PU.1) and SPIB^76–78^ in dendritic cells. We also identified TRs that are broadly functional in multiple PBMC cell types, such as RELA, RELB NFKB2, and RUNX1^79,80^. These findings demonstrate the effectiveness of BARTsc bimodal mode in uncovering lineage-specific functional TRs from single-cell multiome data.

**Figure 3.**
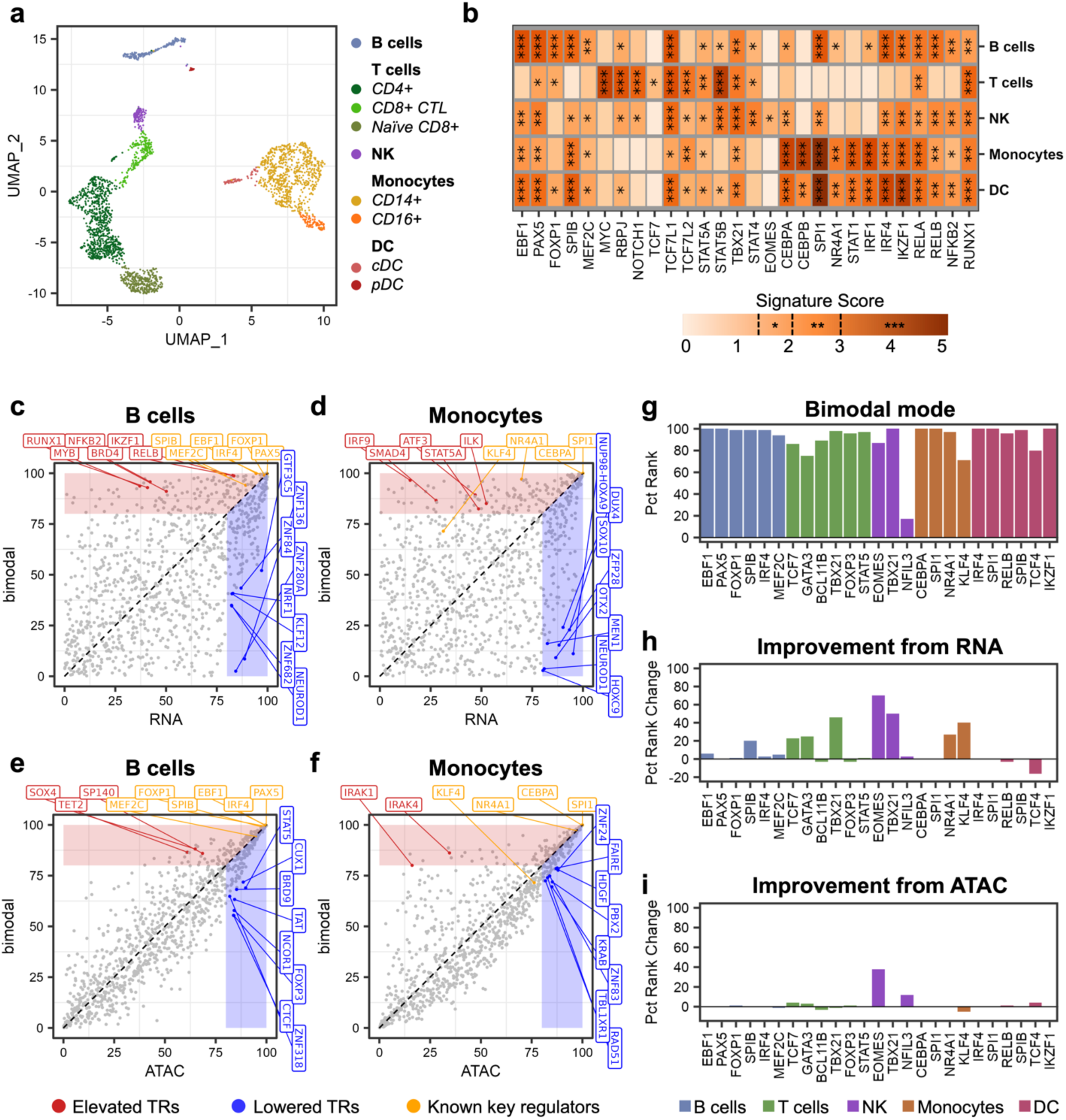
Integrative analysis of scMultiome data improves prediction accuracy. (**a**) UMAP visualization of the scMultiome dataset for human PBMC, where each point represents a single cell colored by cell type. (**b**) Signature analysis results for selected TRs in human PBMC. The heat map is colored by signature score. Asterisks indicate significance level, *: *p* < 0.05, **: *p* < 0.01, ***: *p* < 0.001. (**c**-**f**) Scatter plots comparing the percentile rank (PR) of TRs between bimodal mode and RNA mode (**c**, **d**) or between bimodal mode and ATAC mode (**e**, **f**). Red and blue shaded regions denote the thresholds used to define elevated TRs (c, d: bimodal PR > 80 and bimodal PR > RNA PR; e, f: bimodal PR > 80 and bimodal PR > ATAC PR) and lowered TRs (**c**, **d**: RNA PR > 80 and RNA PR > bimodal PR; **e**, **f**: ATAC PR > 80 and ATAC PR > bimodal PR), respectively. Known key regulators, representative elevated TRs and representative lowered TRs are labelled in different colors. (**g**-**i**) Bar plots showing PR of known key regulators in the bimodal mode (**g**), the increase in PR from RNA mode to bimodal mode (**h**), and the increase in PR from ATAC mode to bimodal mode (**i**). Bars are colored according to the cell type in which the regulator is functionally associated.

To visualize the difference in performance between the bimodal mode and unimodal modes, we next applied BARTsc’s unimodal modes on the same dataset using either RNA or ATAC modality only, and compared the ranks of output TRs with the bimodal mode. Taking B cells and monocytes as examples, we first observed that most known key regulators are ranked high for the corresponding cell types in all three modes (Fig. 3c-f), demonstrating the robustness of BARTsc. For the given cell types, the ranks of some functional TRs were significantly elevated in the bimodal mode than in either RNA or ATAC mode. Conversely, all TRs with the greatest rank reductions either had cell-type-specific functions in non-corresponding cell types or possessed non-specific function in all immune cells, suggesting that our integration methods successfully removed noise by capturing the consistency between the regulatory profiles derived from both modalities via rank aggregation. We also found the TR ranks produced by the bimodal mode exhibit a higher correlation with those from the ATAC mode than from the RNA mode. This is possibly because the regulatory profile used for integration constitutes a larger proportion of the scored UDHSs in the ATAC-derived regulatory profile compared to the RNA-derived regulatory profile. Since chromatin accessibility is known to be related to TR binding activity in a more direct way than RNA expression, this preference towards ATAC modality was not unexpected.

To evaluate the performance of different modes quantitatively, we collected a set of key regulators essential for cell type development or the maintenance of cell identity and examined their percentile rank in the BARTsc output of different modes (Fig. 3g-i). For both RNA and ATAC modes, most key regulators were predicted with a percentile rank higher than 80 (16/25 for RNA mode and 20/25 for ATAC mode, Fig. S3), demonstrating that using either modality alone can already predict most key regulators. However, for certain cell types like NK cells, only a very limited number of key regulators were predicted by either the RNA mode or ATAC mode. The bimodal mode identified both EOMES and TBX21, the two T-box TFs that control NK cell development and the key regulators ranks were generally increased. As a result, 21 out of 25 reached over 80 percentile rank (Fig. 3g). Therefore, we concluded that the bimodal mode improved prediction performance by integrating paired transcriptome and chromatin accessibility modalities.

### Cross-cell-cluster analysis reveals differential TR activities across cell types

We next designed a cross-cell-cluster analysis to assess whether TRs exhibit different functional activities and to quantify such relative activities across different cell clusters. While BARTsc signature analysis can identify TRs that regulate signature gene expression and/or bind to signature open chromatin regions for each cell cluster, it does not differentiate TRs’ relative activity across different cell clusters. Some TRs can be essential for a specific cell type but also function in another, so it is not uncommon that some TRs show high signature scores across multiple cell types (Fig. 2b, 3b). To quantitatively determine whether a TR is relatively more active in one cell cluster over another, we employed a comparative analysis across all possible pairs of cell clusters within the single-cell dataset. We defined a Deviation Ratio (DR) ranging from -1 and 1 to quantify a TR’s activity difference between two cell clusters. For each TR, BARTsc generates a DR matrix capturing all pairwise comparisons between clusters (Fig. 4a). In the DR matrix, a positive DR element indicates a higher TR activity in the row cell cluster relative to the column cell cluster, whereas a negative value indicates higher activity in the column cell cluster relative to the row cell cluster. By taking the row average, BARTsc then summarizes the comparisons between a row cell cluster and all other cell clusters as one value, Mean Deviation Ratio (MDR), indicating the overall relative activity of the row cell cluster (Fig. 4a).

**Figure 4.**
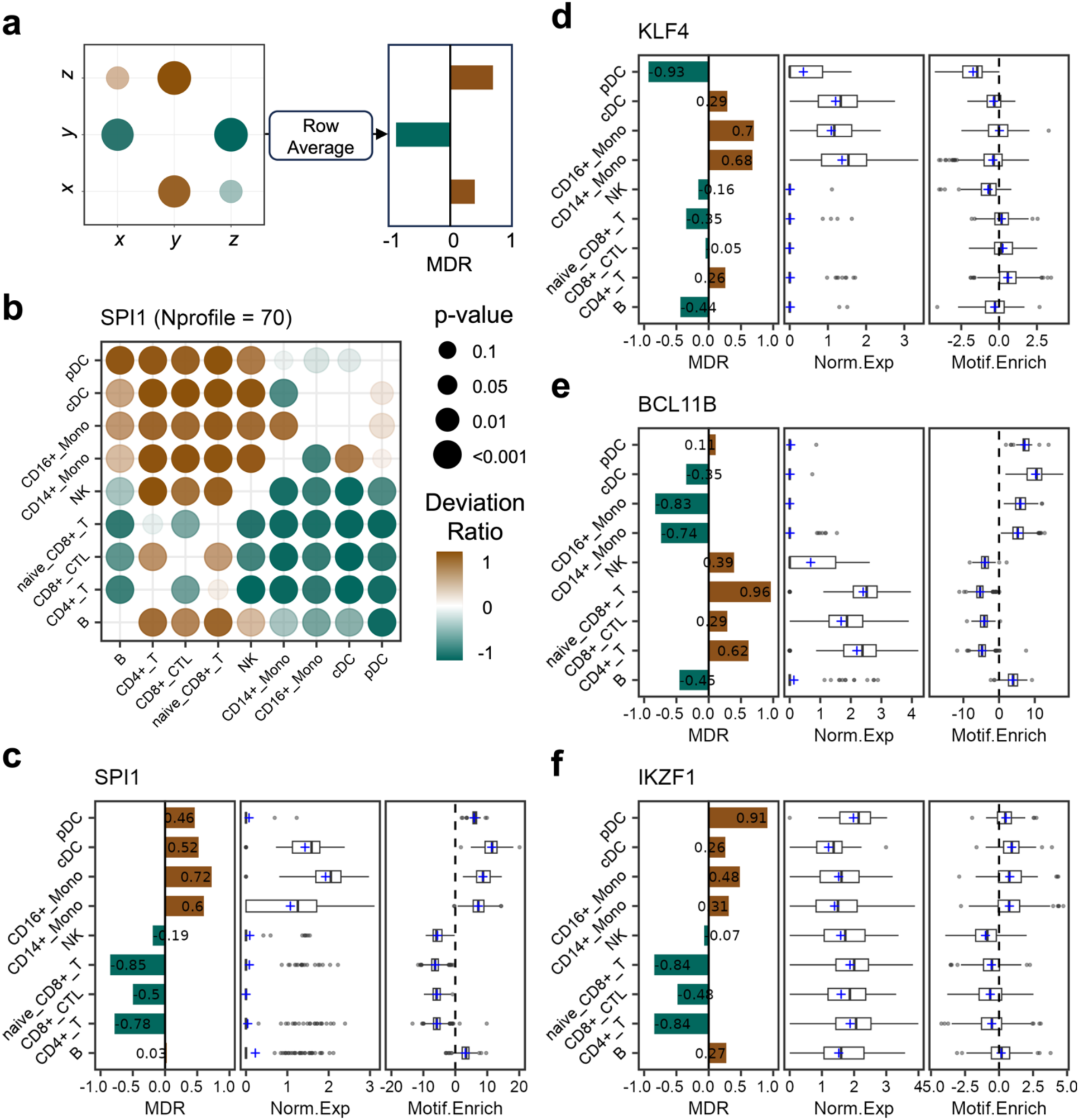
BARTsc reveals relative TR activity across cell types. (**a**) Schematic illustration of DR matrix bubble chart and MDR derived from the DR matrix. (**b**) DR matrix for SPI1. Bubble size indicates the significance level of the pairwise comparison between cell clusters, with larger bubbles corresponding to smaller *p*-values. (**c**-**f**) Relative TR activity metrics for SPI1 (**c**), BCL11B (**d**), IKZF1 (**e**), and KLF4 (**f**). In each panel, Left: MDRs of the TR across PBMC cell types. Middle: distributions of normalized expression level of the TR across PBMC cell types. Right: distributions of the TR motif enrichment scores (chromVAR z-score) across PBMC cell types. In the box plots, center lines represent medians; boxes denote the interquartile range (IQR); whiskers extending to the minimum and maximum values within 1.5× IQR from the median; and blue crosses representing mean values.

To validate whether cross-cell-cluster analysis infers relative TR activity correctly, we applied it to the human PBMC scMultiome dataset. We first examined the DR matrix of SPI1 (PU.1), a master TF for the myeloid lineage^81^. The DR matrix suggests higher SPI1 activity in all myeloid cell types including CD14+ monocytes, CD16+ monocytes, conventional DCs and plasmacytoid DCs compared to any lymphoid cell type (Fig. 4b). Among the lymphoid cell types, SPI1 shows the highest relative activity in B cells, consistent with SPI1’s regulatory role in B cell differentiation and B cell receptor (BCR) signaling^82^. Moreover, The MDRs calculated from this DR matrix is consistent with the pattern of SPI1 gene expression and the pattern of motif enrichment score calculated by chromVAR^83^ (Fig. 4c). These results together with other examples (Fig. S4) confirm the robustness of BARTsc in predicting correct relative TR activities across multiple cell types.

It is known that TR’s transcription level or motif enrichment in open chromatin alone is not always sufficient to infer TR activity, as TRs can function at very low expression level and chromatin accessibility alone does not guarantee TR binding. Remarkably, BARTsc can mitigate these limitations and successfully infer TR activities even in cases where they are not consistently correlated with both TR expression and motif enrichment. For example, KLF4 is a key regulator of monocytes, controlling genes involved in activation and inflammation^84^. In BARTsc results, KLF4 has high MDR scores in monocytes, a feature reflected by KLF4’s expression but not its motif enrichment (Fig. 4d). Similarly, BARTsc predicted high BCL11B activity in T and NK cells, consistent with its known role in maintaining T cell identity and promoting NK cell differentiation^85,86^. However, such patterns were not observed in motif enrichment (Fig. 4e). IKZF1, an important regulator for B cell and pDC development^87,88^, is broadly expressed across all PBMC cell types, with relatively lower expression in B cells and weak motif enrichment in pDCs, making a difficult case for predicting its functional activity.

However, BARTsc identified the highest IKZF1 activity in pDCs and elevated activity in B cells compared to other lymphoid populations, aligning more closely with known functions (Fig. 4f). These examples demonstrated that BARTsc’s cross-cell-cluster analysis under the bimodal mode can more accurately predict TR activity across cell types, overcoming the limitations of relying on gene expression or motif enrichment alone.

### BARTsc outperforms existing methods in identifying key regulators from single-cell data

The ultimate goal of BARTsc is to identify key regulators that play essential roles in establishing or maintaining the identity of each cell type/cluster from a single-cell dataset. To achieve this, BARTsc integrates the cell-cluster-centric perspective derived from signature analysis and the TR-centric perspective derived from cross-cell-cluster analysis to make a comprehensive and accurate prediction of key regulators. We integrated three factors in the process: (1) signature score to quantify the contribution of TRs to the signature expression of the target cell cluster, (2) MDR to quantify the relative TR activity within the target cell cluster compared to other cell clusters, and (3) differential MDR (dMDR) to quantify how unique the TR activity is in the target cell cluster in the examined cell system (see Methods for details). BARTsc uses rank integration for these three factors for each cell cluster and selects top-ranked TRs as key regulators (Fig. 5a-c).

**Figure 5.**
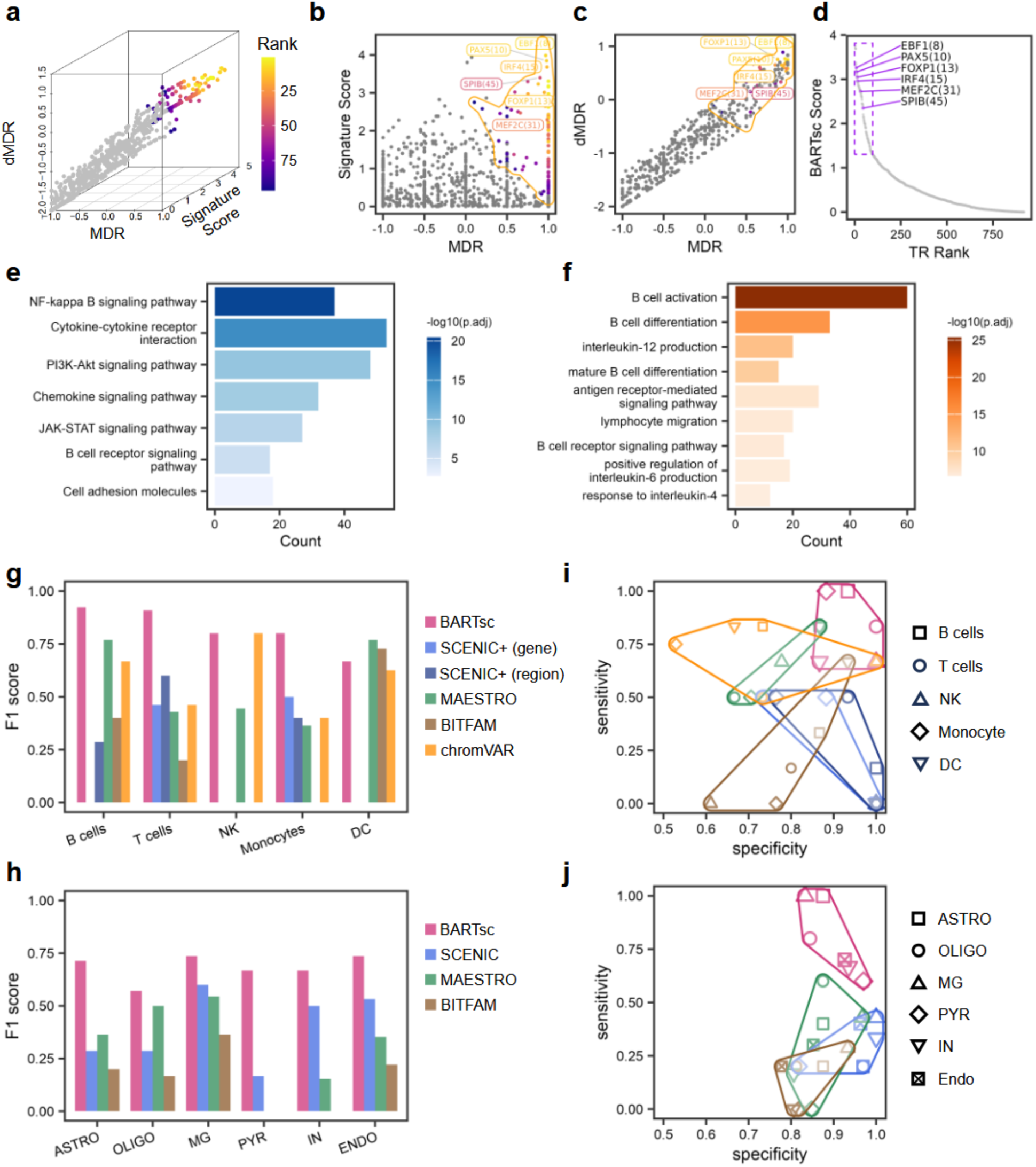
BARTsc achieves highest accuracy in identifying key regulators. (**a**) 3D scatter plot showing TRs positioned based on the three metrics used for B cell key regulator identification. Each dot represents a TR, colored by its integrated rank generated by BARTsc. Grey dots indicate non-key-regulator TRs. (**b**, **c**) Scatter plots showing the relationship between Signature Score and MDR (**b**), and the relationship between dMDR and MDR (**c**). Dots are colored similarly as in (**a**). Known B cell key regulators are labelled with their final rank. (**d**) Scatter plot showing the relationship between BARTsc score and TR final rank for B cell key regulator identification. The purple box indicates BARTsc-predicted key regulators with BARTsc scores above the default cutoff. Known B cell key regulators are labelled with their final rank in the parenthesis. (**e**, **f**) Selected KEGG pathways (**e**) and Gene Ontology terms (**f**) enriched for BARTsc-predicted B cell key regulators and their target genes. (**g**-**j**) Performance comparison between BARTsc and other methods in identifying key regulators for each cell type from human PBMC scMultiome dataset (**g**, **i**) and mouse cortex scRNA-seq dataset (**h**, **j**). Bar plots show the F1 scores for key regulator prediction for different methods across different cell types in human PBMC (**g**) and mouse cortex (**h**). Scatter plots show the sensitivity and specificity of each method in identifying cell type key regulators in human PBMC (**i**), and mouse cortex (**j**).

Taking B cells in the PBMC dataset as an example, we were able to identify all the known key regulators for B cells ranked among the highest in the BARTsc result (Fig. 5d). EBF1 and PAX5 were ranked among the top 10, consistent with their core role in activating B-cell-specific gene expression and repressing genes associated with other lineages.^62,63^ Other BARTsc-predicted top TRs including BACH2, RELB, NCOR2, POU2F2 and CD74 are also known for their direct participation in B cell development or function^89–93^ (Fig. S5a). Our functional analysis on the BARTsc-predicted key regulators and their target genes showed significant enrichment of KEGG Pathways essential for B cell differentiation, proliferation and activation, such as NF-kappa B signaling pathway and B cell receptor signaling pathway (Fig. 5e), and Gene Ontology terms related to B cell functions (Fig. 5f). We also identified key regulators and pathways critical for the development and functions for other PBMC cell types as well (Fig. S5b-e, Fig. S6). Similarly, we showed that BARTsc identified almost all the known key regulators as well as other widely mentioned functional TRs for each cell type from the mouse cortex dataset (Fig. S5f-k). Taking together, our results demonstrated that BARTsc can accurately identify key regulators supported by existing knowledge and regulate genes associated with cell-type-specific functions.

To systematically assess the performance of BARTsc in comparison with other existing methods in identifying key regulators, we used the human PBMC scMultiome dataset and the mouse cortex scRNA-seq dataset to evaluate each method’s accuracy to identify known key regulators for each cell type. Other existing methods to compare include SCENIC+^94^, MAESTRO^27^, BITFAM^26^ and chromVAR^83^, all of which are representative tools for similar tasks. Our results showed that BARTsc achieved the highest F1 score compared to other methods in four out of five human PBMC cell types (Fig. 5g) and all six cell types in mouse cortex (Fig. 5h). Notably, in human T cells and monocytes, BARTsc outperformed SCENIC+, the second-best method, by a substantial margin. In addition, BARTsc achieved both higher sensitivity and specificity than other methods while maintaining a balance between them (Fig. 5i, j). These results highlight the robust and superior performance of BARTsc in accurately identifying key regulators across diverse cell types compared to other methods.

### BARTsc identified NELFA, a novel proliferation driver in pancreatic cancer

TRs play central roles in cancer initiation, progression, and heterogeneity, making their study critical for understanding cancer biology and identifying potential therapeutic targets. To evaluate BARTsc’s ability to identify functional TRs in cancer, we re-analyzed a single-cell Multiome dataset of pancreatic ductal adenocarcinoma (PDAC) (Fig. 6a). Using BARTsc, we successfully identified established key regulators across all major cell types, spanning a wide range of cell populations from secretory cell types (acinar cells, ductal cells, endocrine cells) to supportive cell types (endothelial cells, fibroblasts, myeloid cells, T cells) (Fig. 6b). Similar to the results on other datasets, BARTsc outperformed existing methods in accurately identifying key regulators in most PDAC cell types (Fig 6c, d), and can uniquely identify several known key regulators that are missed in all other methods, such as TAZ for acinar-to-ductal metaplasia (ADM)^95^ and NR4A1 for myeloid cells^96^. Focusing on the cells undergoing ADM, a critical event in PDAC initiation, we identified not only established marker TRs for both acinar cells (HNF1B^97^) and ductal cells (KLF4^98^), but also SOX9, a TF with an indispensable role for ADM induction^99^ (Fig. 6b). Functional analysis of BARTsc-identified key regulators and their putative target genes highlighted pathways that play significant roles during ADM, such as Hippo, PI3K-Akt, MAPK signaling pathways^100^ and morphogenesis-related Gene Ontology terms^101^ (Fig. S7a, b). Furthermore, for ADM cells, SOX9 ranked higher in BARTsc than any other methods among not only all TRs, but also within the SOX family members sharing similar TFBMs (Fig. S7c, d). These results further demonstrated BARTsc’s superior performance in accurately identifying key regulators from cancer single-cell data.

**Figure 6.**
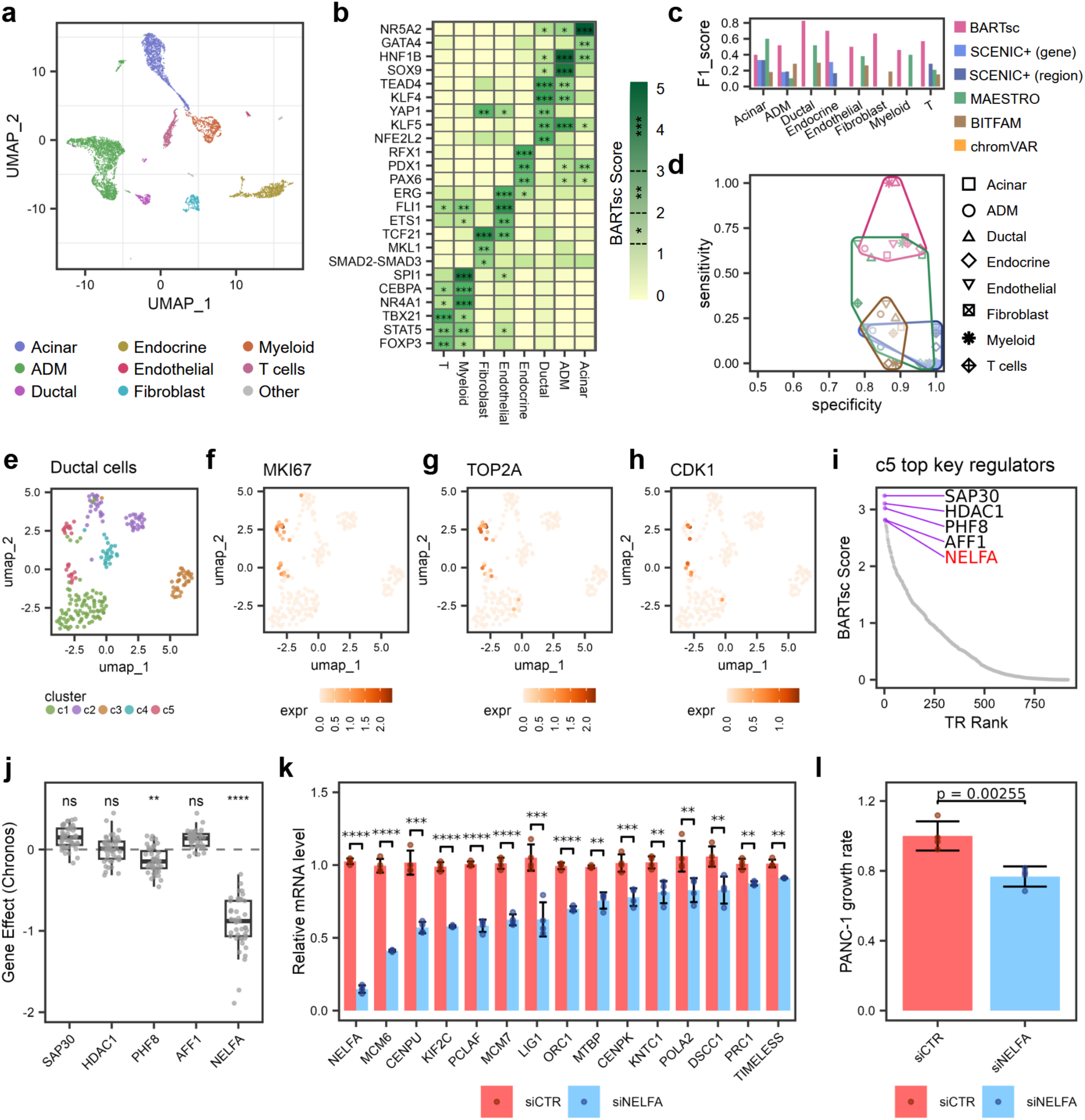
BARTsc identifies functional TRs in pancreatic cancer (PDAC). (**a**) UMAP visualization of the scMultiome dataset for human PDAC, where each point represents a single cell colored by cell type. (**b**) Signature analysis results for selected TRs in human PDAC. The heat map is colored by BARTsc score. Asterisks indicate significance level, *: *p* < 0.05, **: *p* < 0.01, ***: *p* < 0.001. (**c**, **d**) Performance comparison between BARTsc and other methods in identifying key regulators for each cell type from PDAC scMultiome dataset. Bar plots show the F1 scores for key regulator prediction for different methods across different cell types (**c**). Scatter plot shows the sensitivity and specificity of each method in identifying cell type key regulators (**d**). (**e**-**h**) UMAP visualizations of the ductal cells from the PDAC scMultiome dataset, where each point represents a single ductal cell colored by sub-clusters (**e**), or the normalized expression level of gene MKI67 (**f**), TOP2A (**g**) and CDK1 (**h**). (**i**) Scatter plot showing TRs ranked by BARTsc score. The top five predicted key regulators are highlighted and labeled. (**j**) Chronos scores quantifying the effects of CRISPR-mediated knockout of the top five predicted key regulators in PDAC cell lines. Each point represents a PDAC cell line, with lower Chronos scores indicating greater loss of cell fitness upon gene knockout. P-values are calculated by one-sided Wilcoxon rank-sum tests. (**k**) RT-qPCR results showing the relative mRNA levels of NELFA and selected genes upon NELFA knock-down by siRNA in PANC-1 cells. Bars represent the mean mRNA levels relative to those under control siRNA transfection groups after normalization to *GAPDH* mRNA levels, and error bars indicate the standard deviation. Asterisks indicate significance levels between siNELFA and siControl determined by one-sided *t*-test, *: *p* < 0.05, **: *p* < 0.01, ***: *p* < 0.001. (**l**) Growth of PANC-1 cells three days after NELFA knockdown (siNELFA) compared with the control cells (siControl). Error bars indicate the standard deviation. P-value is calculated by one-sided *t*-test.

PDAC, by definition, is composed of malignant epithelial cells that adopt a ductal-like identity; however, these ductal-like tumor cells are not homogeneous, and substantial intratumoral heterogeneity exists across ductal malignant programs and states^102,103^. Therefore, we further clustered the ductal cells into 5 distinct sub-clusters (Fig. 6e) and identified one sub-cluster (c5) as particularly notable for its high expression of proliferation markers (Fig. 6f-h) and reduced expression of tight junction-related genes, suggesting a rapidly growing cancer cell state and the loss of epithelial polarity^104^ (Fig. S8a-c). Remarkably, we found that expression of c5 signature genes is significantly associated with poor prognosis in The Cancer Genome Atlas (TCGA) data (Fig. S8d), and that these genes were enriched for multiple cell cycle-related pathways such as cell division and DNA replication (Fig. S8e). Additionally, we found these cells’ low differentiated level and high potency level estimated using CytoTRACE2^105^ (Fig. S8f, g), suggesting that this cell sub-cluster represented a highly proliferative and aggressive tumor cell population. We next applied BARTsc to the ductal cell sub-dataset to identify candidate key regulators, which might play critical roles in PDAC proliferation. The top TRs for c5 predicted by BARTsc include SAP30, HDAC1, PHF8, AFF1 and NELFA (Fig. 6i), all of which have been implicated in cancer. By examining the CRISPR screening data from DepMap^106^, we found that the perturbation of NELFA, but not the other candidate TRs significantly impaired proliferation of PDAC cells (Fig. 6j). Based on these observations, we predicted that NELFA is a key regulator of the high proliferative PDAC cell population. To validate this finding, we performed RNAi-mediated knockdown of NELFA in the PANC-1 cell line (Fig. S8h, i) and found that c5 signature genes involved in cell division (e.g., CENPU, KIF2C) or DNA replication (e.g., MCM6, MCM7) were significantly downregulated upon NELFA knockdown (Fig. 6k), suggesting the upstream regulatory function of NELFA. Consistently, we also found that NELFA knockdown significantly retarded the growth of PANC-1 (Fig. 6l). Taken together, with experimental validation, we identified NELFA as a novel driver of PDAC proliferation, acting by regulating cell-division-related genes. These data further demonstrated the power of BARTsc in identifying novel functional regulators from single-cell data.

## Discussion

Investigating functional TRs is essential for understanding transcriptional regulation mechanisms underlying heterogeneous cell systems. The rapid development of single-cell sequencing techniques now enables predicting functional TRs at the cell type or sub-type levels from transcriptomic or chromatin accessibility profiling data. As presented in this work, BARTsc is a method that directly uses public ChIP-seq data as TR binding reference and leverages differential genomic features between cell states in single-cell data to predict functional TRs.

BARTsc takes advantage of the unique features from both single-cell data and public ChIP-seq data and integrates them for accurate TR prediction. A major advantage of single-cell sequencing compared to bulk sequencing is its ability to resolve cellular heterogeneity, thus various cell types or cell states within a single-cell dataset can form contrast for each other and be leveraged to identify their differential genomic features. In doing so, BARTsc not only can analyze how TRs regulate cell-type-specific features but also can uncover differential activity of TRs between any pair of cell clusters. Meanwhile, as ChIP-seq is the direct measurement of genome-wide TR activity, BARTsc achieves high-performance TR prediction by associating differential genomic features from single-cell data directly with TR binding profiles characterized by ChIP-seq, an approach used to quantify TRs’ contribution to cell-type signatures and differential TR activity across cell types/clusters.

We evaluated the performance of BARTsc on both scRNA-seq data and single-cell multiome data, and demonstrated that BARTsc can accurately predict functional TRs from datasets across diverse cell systems and species. The bimodal mode of BARTsc significantly enhances prediction accuracy from unimodal mode by integrating information from both scRNA-seq and scATAC-seq. In practice, this approach can be applied for analyzing both matched-modal single-cell data (multiome with RNA and ATAC co-assayed from the same cell) and unpaired scRNA-seq and scATAC-seq data (assays on separate materials from the same biological sample), thereby extending the usability of BARTsc to a broader range of datasets. The cross-cell-cluster analysis of BARTsc enables an accurate assessment of TR activity across various cell types and subtypes in a complex tissue, outperforming conventional approaches relying solely on gene expression or sequence motif information. Finally, BARTsc enables a comprehensive identification of key regulators by integrating signature analysis and cross-cell-cluster analysis, showing a significantly better performance and short runtime (Fig. S9) compared to existing methods in both scRNA-seq and scMultiome scenarios.

We applied BARTsc to a human PDAC dataset and identified established key regulators across this highly heterogeneous tumor context and uncovered NELFA as a previously unrecognized regulator of pancreatic tumor proliferation. As a core component of the negative elongation factor (NELF) complex, NELFA is essential for promoter-proximal pausing of RNA-polymerase II (Pol II)^107,108^. By facilitating the maintenance of promoter-proximal chromatin accessibility and integrating multiple regulatory signals, Pol II pausing functions as a regulatory mechanism distinct from Pol II recruitment and elongation^109–111^. Previous studies have shown that loss of NELF impairs the development of colon cancer and breast cancer, underscoring its importance in the maintenance of cancer stemness^112,113^. Our experiments characterized NELFA’s pro-tumor function in PDAC and further discovered its specific activity in a high-proliferative subset of pancreatic tumor cells. Our experimental data established a regulatory link between NELFA and cell cycle-related genes, providing mechanistic insights into its essential role in tumor cell proliferation and highlighting NELFA as a potential candidate of therapeutic targets and for further investigation.

Several limitations are present with this method. First, the range of TRs that can be predicted by BARTsc is limited by existing ChIP-seq data. Although the current ChIP-seq data collection has included nearly 1000 TRs for human and over 500 TRs for mouse, there are still some TRs that do not have high-quality ChIP-seq data available, likely due to the lack of high-quality antibodies. Second, while the BART algorithm is designed to integrate multiple ChIP-seq datasets from various cell types for the same TR to increase the prediction power and reduce biases, such potential cell type selection biases may still exist, especially for TRs with few ChIP-seq datasets. For example, there is only one NFIL3 ChIP-seq dataset in our data collection, which may explain why BARTsc failed to identify NFIL3 as an active TR in NK cells. Third, BARTsc is designed and evaluated on clustered single-cell data, which carries a strong assumption that cell clusters are distinct in a relatively discrete way. However, some single-cell datasets may be better characterized by continuous trajectories reflecting gradual cell state transitions. While we believe that BARTsc can still identify key transcriptional regulators that drive the continuous cell state transition by leveraging the differential features (RNA or ATAC) along the continuous trajectory, we have not yet conducted a systematic evaluation in this context.

Despite these limitations, BARTsc represents a significant methodological advance in transcriptional regulation analysis of single-cell genomics data. The innovative approach of integrating cell-cluster-centric information and TR-centric information to comprehensively determine key transcriptional regulators guarantees BARTsc a superior performance than existing methods. Implemented as an open-source package, BARTsc is fully accessible to the community as a valuable tool for elucidating cell-type-specific regulatory programs and facilitating deeper insights into transcriptional regulation across diverse biological systems.

## Methods

### BARTsc algorithm Data preprocessing

BART is designed to analyze a single-cell omics dataset in which cells have been grouped into clusters or cell types. The input of BARTsc includes the count matrix and cell type/cluster labels for each cell. BARTsc first performs data normalization according to the input data type. Specifically, BARTsc performs log-transformed library-size normalization on the cell-by-gene matrix for scRNA-seq, and term frequency-inverse document frequency (TF-IDF) normalization on the cell-by-peak matrix for scATAC-seq^114^. Using the normalized expression (or accessibility) matrix, BARTsc next looks for cell-cluster-signature features and pairwise differential features by comparing each cell cluster with all other cells and comparing each pair of cell clusters, respectively. These two types of features sets are collectively referred to as differential feature sets. BARTsc uses presto^115^ to perform Wilcoxon rank-sum test and Benjamini-Hochberg adjusted P-value for differential feature selection.

### Inference of cis-regulatory profile

BARTsc uses union DNaseI hypersensitive sites (UDHS) as a repertoire of all potential cis-regulatory regions for genome-wide profiles^38^. Each differential feature set is used to predict a cis-regulatory profile, which consists of regulatory scores for each DHS. BARTsc processes differential feature sets differently depending on input data type. For scRNA-seq, each differential feature set is a gene set. BARTsc uses a compendium of H3K27ac profiles (7968 human datasets and 5851 mouse datasets) to quantify cis-regulatory activities. Each H3K27ac profile can be represented by a gene activity profile generated by calculating the regulatory potential on each gene as described^38^. BARTsc conducts adaptive lasso regression to find a weighted combination of selected H3K27ac gene activity profiles that best matches the input gene set, which is represented by a binary vector across all the genes. The vector sum of the selected H3K27ac profiles weighted by their regression coefficients is then used as the cis-regulatory profile of the given gene set. For scATAC-seq, each differential feature set is a peak set scored by log2 scaled fold change. Since differentially accessible regions and their fold change signals intuitively reflect the differential cis-regulatory elements and the change of their regulatory activity, respectively, BARTsc directly maps the scored peak set to UDHSs as the cis-regulatory profile. For single-cell multiome, the differential feature sets, including a gene set and a scored peak set derived from the comparison between the same pair of cell clusters, are used together to predict a cis-regulatory profile. BARTsc first predicts a gene set-derived cis-regulatory profile and region set-derived cis-regulatory profile using the two parallel differential feature sets individually and then integrate them as a consensus cis-regulatory profile using geometric mean-based rank aggregation. A typical differentially accessible peak set usually covers only a small proportion of UDHS, resulting in a sparse region set-derived cis-regulatory profile (with many zero-scored sites) compared to a gene set-derived cis-regulatory profile. To avoid an imbalanced consensus cis-regulatory profile biased towards the RNA modality, BARTsc uses a maximum slope-based strategy to find the optimal number of top sites to consider and ignores the rest. In brief, BARTsc first sorts both cis-regulatory profiles by regulatory score and then for every top x selected sites, counts y, the number of sites that are included in the top x sites for both profiles. Subsequently, BARTsc uses generalized additive model to fit the relationship between x and y as function y = f(x) and uses x_max_ that leads to the maximum slope as the optimal number of top sites. For both profiles, only top x_max_ UDHSs are considered in rank aggregation so that scRNA-seq and scATAC-seq contribute to the consensus cis-regulatory profile in a balanced way.

### Calculation of association score

To evaluate the association between cis-regulatory profile and a given ChIP-seq dataset, we first map the ChIP-seq peaks onto UDHS to generate a binary binding profile indicating whether each UDHS overlaps with the peaks or not. Then, BARTsc sorts the cis-regulatory profile by regulatory score and uses receiver-operating characteristic (ROC) curve to evaluate the association between the given cis-regulatory profile and each TR binding profile. The area under the ROC curve (AUC) value is calculated for each TR binding profile and is termed as association score.

### Cell cluster signature analysis

BARTsc uses cis-regulatory profiles derived from cell type signatures to estimate the contribution of TRs to cell-type signature features. For each signature feature set, BARTsc first performs cis-regulatory profile inference and association score calculation to generate an association profile. Then, BARTsc performs one-tailed Wilcoxon rank sum test between the association scores of each given TR’s binding profiles and those of all the rest binding profiles. Since the distributions of TR binding sites greatly vary, some TRs’ binding profiles are more likely to receive high association scores than others. A standardization step to minimize this inequality between TRs is thus in need. Our strategy is to first perform signature analysis on a compendium of representative inputs, and then use their outputs as a baseline to correct future predictions. We collected the gene sets from the Molecular Signatures Database (MSigDB) with at least 200 genes as gene set compendium and H3K27ac profiles as chromatin profile compendium for scATAC-seq. For each TR 𝑖, we calculate a standard Z-score:

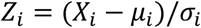

where X_i_ is the Wilcoxon statistic score of TR 𝑖 calculated from the query data, 𝜇_i_ and 𝜎_i_ are mean and standard deviation of Wilcoxon statistic scores of TR 𝑖 across all datasets from the compendium. To give a summarized result, BARTsc first ranks TRs by standard Z-score, Wilcoxon P value and maximum AUC value individually and takes the average rank as the final rank of TRs. A final P value is then calculated for each TR based on Irvin-hall distribution as background. In this final output, the higher a TR rank is, the more likely that TR is responsible for the expression of the signature gene set.

### Cross-cell-cluster analysis

To investigate TR binding activity across cell clusters, for each pair of cell clusters, BARTsc first identifies a pair of differential feature sets by comparing the given cell cluster with each other. BARTsc uses each differential feature set for cis-regulatory profile prediction and association score calculation, so that a pair of ChIP-seq association profiles will be generated. Next, for each TR, BARTsc performs two-tailed Wilcoxon signed test on the paired ChIP-seq association scores of the given TR’s ChIP-seq samples to test whether the given TR is differentially associated with the two cell types under investigation. To quantify the association difference, BARTsc calculates a Deviation Ratio for each TR and for all the possible comparisons between any two cell clusters *x* and *y*:

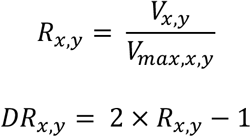

where *V* is the *V* statistics of Wilcoxon signed test and *V_max_* is the maximum possible value of *V* statistics from the test. *R* is a normalized ratio of V statistics. Then, *R* is rescaled to the range from -1 to 1, denoted as *Deviation Ratio* (*DR*), which represents the extent to which the given TR is more associated with one of the two cell types being compared. Specifically, the closer *DR* is to 1, the more associated the TR is to cell type *x*, and the closer *DR* is to -1, the more associated the TR is to cell type *y*. Finally, for each TR, An *N* × *N DR* matrix is generated, for a given dataset with *N* cell clusters and each entry represents the comparison between the row cell cluster and the column cell cluster. The row average of the DR matrix can be further taken to simplify the output, resulting in a *Mean Deviation Ratio* (*MDR*) value for each cell cluster.

### Identification of key regulator for each cell cluster

BARTsc identifies cell cluster’s key regulators by summarizing signature analysis and cross-cell-cluster analysis. For a given TR, we quantify the uniqueness level of its activity in cell cluster *x* as follows:

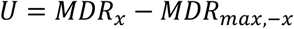

Where *MDR_max,-x_* denotes the maximum *MDR* among all other cell clusters, which is subtracted from the *MDR* of cell type *x* as the measurement of uniqueness, and *U* denotes the uniqueness. BARTsc first ranks TRs by signature score, MDR and uniqueness score individually and then aggregates these ranks in order to generate a consensus importance ranking of TRs. Since rankings follow uniform distribution, BARTsc uses Irvin-hall distribution to estimate the p values of the consensus ranking:

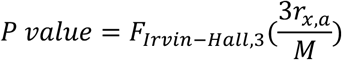

Where *F_Irwin-Hall,3(⋅)_* represents the cumulative distribution function (CDF) of the Irvin-hall distribution with n=3, while *r_x,a_* is the final importance rank of TR *a* for cell cluster *x* and *M* is the total number of TRs. In this consensus TR ranking, the higher the TR rank is, the more likely that TR is a key regulator for cell cluster *x*.

### Re-analysis of published data

#### Mouse cortex scRNA-seq dataset

The raw count matrix and corresponding cell type annotations of the mouse cortex scRNA-seq dataset were retrieved from the official website associated with the publication^39^ (see code and data availability). Seurat^116^ (version 4.0.5) was used to perform Principal Component Analysis (PCA) and to calculate Uniform Manifold Approximation and Projection (UMAP) for visualization with the top 14 principal components (PCs).

#### Human PBMC scMultiome dataset

Processed data of the PBMC dataset was retrieved from 10X Genomics website (see code and data availability). We used Seurat^116^ (version 4.0.5) and ArchR^117^ (version 1.0.2) for the quality control of the RNA modality and ATAC modality respectively. Barcodes with RNA count were filtered by nFeature_RNA > 200 & nFeature_RNA < 4000 & percent.mt < 20. Barcodes with ATAC count were filtered by fragment number > 1000 and TSS enrichment > 4. Function addDoubletScores from ArchR was then used to infer doublets so that only singlets were kept. Barcodes passed both RNA-based QC and ATAC-based QC were kept as high-quality cells with double-modality. Clustering and visualization were performed with Seurat based on the RNA modality as follows: We first identified the top 2000 highly variable genes and performed PCA. Next, we selected the top 13 PCs to construct shared nearest neighbor graph and conducted clustering using Louvain algorithm with a resolution of 0.5. We used the same PCs to generate UMAP. We annotated cell types based on the expression of known marker genes (Fig. S2).

#### Human PDAC scMultiome dataset

We first used 10X Genomics Cell Ranger Arc^118,119^ (version 2.0.2) to process the FASTQ files of the PDAC dataset, and then used Seurat to filter cells based on the thresholds used by the publication^120^. We used Seurat for clustering and visualization based on the RNA modality. Similar to the analysis of the PBMC dataset, we identified the top 2000 highly variable genes for PCA. We next removed batch effect using Harmony^121^ (version 1.2.3) and used the top 20 harmony-corrected PCs to construct shared nearest neighbor graph and generated the UMAP visualization. We clustered all the cells using the Louvain algorithm with a resolution of 0.2 first to separate cell types roughly. We labeled the cells using the cell type annotation in the original publication. We then subset the cell clusters annotated as ductal cells for sub-clustering. We performed PCA and harmony correction on the subset of cells again and used the top 20 harmony-corrected PCs to construct shared nearest neighbor graph followed by a Louvain clustering with a resolution of 0.5.

#### Functional enrichment analysis

To assess the functional pathways regulated by BARTsc-predicted key regulators, for a given cell type, we compiled a combined list of predicted key regulators and their target genes, which was then subjected to functional enrichment analysis. TF-target gene interactions with confidence levels equal or higher than level C from DoRothEA gene regulatory network^122^ (Version 1.18.0) were used to determine target genes. Gene Ontology and KEGG functional enrichment analyses were conducted using function enrichGO and enichKEGG from clusterProfiler^123^ (version 4.10.0), respectively.

### Comparison of BARTsc with existing tools

#### Benchmark standards and tool selection

We assessed the performance of BARTsc, along with other tools of similar functionality, by their ability to predict functional transcriptional regulators (TRs) for each cell cluster. To quantify the assessment, we compared the relative ranks assigned to known functional TRs in each tool’s output, under the assumption that the known functional TRs should rank high in a good-performing tool’s output. We also examined each tool’s sensitivity and specificity to identify key regulators for each known cell type. We selected existing bioinformatics tools with functionalities comparable with BARTsc based on the following criteria: 1) the tool is designed for predicting TRs or TR regulons from single-cell omics data, and 2) the output of the tool includes a ranked list of TRs for each cell cluster, enabling direct comparison with BARTsc output. Based on these criteria, we identified SCENIC^17^, MAESTRO^27^ and BITFAM^26^ as comparable tools for analyzing scRNA-seq data or the RNA modality of single-cell multiome data, and SCENIC+^94^ as a comparable tool for single-cell multiome data, as well we ChromVAR^83^ (through ArchR^117^) as a comparable tool for the ATAC modality of single-cell multiome data. We ran all the methods that support parallel computation (BARTsc, SCENIC, SCENIC+, BITFAM, and ChromVAR) with 6 CPU cores under identical hardware settings, to ensure a fair comparison of runtime.

#### AI-assisted collection of ground truth

A major challenge in validating and benchmarking TR prediction tools is the lack of ground-truth knowledge. We need an extensive collection of TR-cell type relationships supported by literature to curate a functional annotation table that documents the functional roles of TRs in specific cell types. To achieve this, we developed and employed an AI-assisted approach to systematically screen peer-reviewed literature to identify and summarize functional TR-cell type relationships with validation.

We implemented this approach as an automatic pipeline, which takes a TR-cell type pair as input and produces either a summary of the TR function in that cell type or a rejection of the proposed relationship as output. We employed the large language model (LLM) GPT-5 mini (OpenAI) for semantic analysis and summary generation. This tool first constructs a search query and retrieves relevant publications from NCBI PubMed^124^. It then iteratively processes each publication through following steps:

1. Download the full text of that publication, go through the full text to extract relevant paragraphs containing both the queried TR and cell type within a specific distance.
2. For each relevant paragraph, use LLM to generate a paragraph to summarize the TR’s functional role in the queried cell type.
3. Combine the paragraph-level summaries into an overall summary for that publication.
4. Reassess the relevant paragraphs with the LLM to determine whether each paragraph supports, rejects, or adds evidence to the overall summary. A self-verification report is generated in this step.
5. Integrate the overall summary and verification report to produce the final output for that publication.

Because the primary objective is to determine whether the queried TR is functional in a specified cell type, the iteration stops once supportive evidence is identified. We then generated the final summary with citations to the original publication. Finally, we manually reviewed the resulting functional annotation table to ensure authenticity.

#### Benchmarking tools for identifying functional TRs

For each cell type, we classified the TRs into 3 categories based on their functional annotations and expression levels: 1) functional: TRs that are expressed and with known function in the given cell type; 2) function-unknown: TRs that are expressed in the given cell type but lack known functional evidence; and 3) non-expressed: TRs that are barely expressed in the given cell type (95th percentile normalized expression < 0.1). Because different tools report varying numbers of TRs in the output, and some report only the TRs that are statistically significant, to ensure a fair comparison, we compared the rank percentiles of the TRs across different tools. For each cell type, we only included the TRs that were present in a tool’s output. We used the median rank percentile of functional TRs as the primary metric to determine the best-performing tool.

#### Benchmarking tools for identifying key regulators

We composed a list of known key regulators for each cell type as the ground truth, based on experimental evidence that the perturbation of a key regulator in a cell type results in severe consequences such as impaired differentiation or loss of cell identity. We used each tool’s default output of cell-type-specific or cell-type-significant TRs for performance evaluation. For BARTsc, we used the output TRs with a final P value < 0.05 in the key regulator identification step. For SCENIC and SCENIC+, we used the output TRs with highest regulon specificity scores (RSS) in a cell type. For MAESTRO, we used the default output of driver TRs. For BITFAM, following the default cutoff, we used the TRs with inferred activity higher than the 75^th^ percentile of all the inferred TRs. For ChromVAR, we applied Wilcoxon rank sum test^115^ to compare the output deviation scores (z-scores) between each cell type and all the other cells and used the TRs with adjusted P value < 0.05 and fold-change > 2 as final output. We then calculate the sensitivity, specificity, and F1 score by comparing the ranked TRs predicted with the ground-truth key regulators to assess the performance of each tool.

### Experimental validation of NELFA as a proliferation driver in PDAC

#### Cell culture

Human pancreatic carcinoma cell line PANC-1 was originally purchased from the American Type Culture Collection (ATCC) (Cat# CRL-1469). They are cultured in complete growth DMEM medium (Dulbecco’s Modified Eagle’s Medium) with 10% fetal bovine serum (FBS) and 1% penicillin-streptomycin (P/S). Cells are incubated at 37℃ with 5% CO_2_ and routinely tested for mycoplasma contamination using Mycoplasma Detection Kit-DigitalTest v2.0 (BioTool, Cat#B39132).

#### Quantitative reverse transcription PCR (RT-qPCR)

Total RNAs were extracted using TRIzol Reagent (Invitrogen) and quantified by Cytation 5 Cell Imaging Multi-Mode Reader (BioTek). 1-2 μg of RNAs were reverse transcribed into cDNA using MultiScribe Reverse Transcriptase (Thermo Fisher Scientific). qPCR reactions were conducted in the PowerTrack SYBR Green Master Mix (Thermo Fisher Scientific) using the QuantStudio3 Real-Time PCR Systems (Thermo Fisher Scientific). Levels of tested transcripts were determined with the ΔΔCt method and normalized to the expression of GAPDH, which serves as the internal control.

#### Protein lysis and Western blot

Total proteins were extracted as previously described^125,126^. Basically, PANC-1 cells were lysed in RIPA buffer (150 mM sodium chloride, 1.0% NP-40, 0.5% sodium deoxycholate, 0.1% SDS, 50 mM Tris pH 8.0) with 1 × protease inhibitor cocktail (Roche) for 1hr at 4℃ with rotating. Protein concentrations were measured using Pierce™ BCA Protein Assay Kits (Thermo Scientific) by Cytation 5 Cell Imaging Multi-Mode Reader (BioTek). Protein lysates were subjected to SDS-PAGE electrophoresis and finally analyzed using Western blots. Antibodies that were used in the immunoblotting include the anti-NELFA Antibody (FORTIS LIFE SCIENCES, Cat#A301-910A) and the secondary anti-rabbit antibody conjugated to HRP (Invitrogen, Cat# A16110). Intensities of immunoblotting bands were quantified using ImageJ.

#### Cell viability and proliferation assays

PANC-1 cells were seeded into 96-well plates and reached 80% confluency the next day. The siRNA targeting either control (siGENOME control pool, Dharmocaon, Cat#D-001206-14-20) or NELFA gene (ON-TARGETplus siRNA, Horizon, Cat# L-012156-00-0005) were transfected into cells using the Lipofectamine® RNAi MAX Reagent (Invitrogen, Cat# 13778100), with a final concentration of 30 nM in each well. 72 hours post-transfection, cell viability was determined using CCK-8 kit (Dojindo, Cat# CK04-01) according to the manual. Briefly, 10 uL of CCK-8 solution was added into each well and incubated at 37℃ for 2 hr. The readings were measured at the absorbance of 450 nm using Cytation 5 Cell Imaging Multi-Mode Reader (BioTek). Cell viability can be calculated with the following formula:

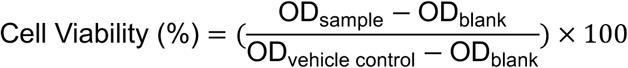

Both control and knockdown groups were repeated with four replicates.

### Statistical analysis

All the experiments were repeated with at least three biological and technical replicates unless otherwise stated. Unpaired Student’s t test was used to assess the statistical significance of the difference between the two groups.

## Code Availability

BARTsc is implemented in R as an open-source package. The package with source code is freely available at https://github.com/zang-lab/BARTsc

## Acknowledgements

The authors thank Todd W. Bauer, Clint L. Miller, Yuh-Hwa Wang, and members of the Zang lab for helpful discussions. H.Z. thanks Emily E. Byrd for help with cell-type annotation and critical review of the manuscript. This work was supported by NIH grants R35GM133712 (C.Z.), R01GM140084 (K.X.), and R01CA279681 (K.X.). H.Z. is a Wagner Fellow in UVA School of Medicine and receives a Farrow Fellowship from the UVA Comprehensive Cancer Center. This work was also partially supported by Saunders funding within the UVA Comprehensive Cancer Center.

---

**Figure S1.**
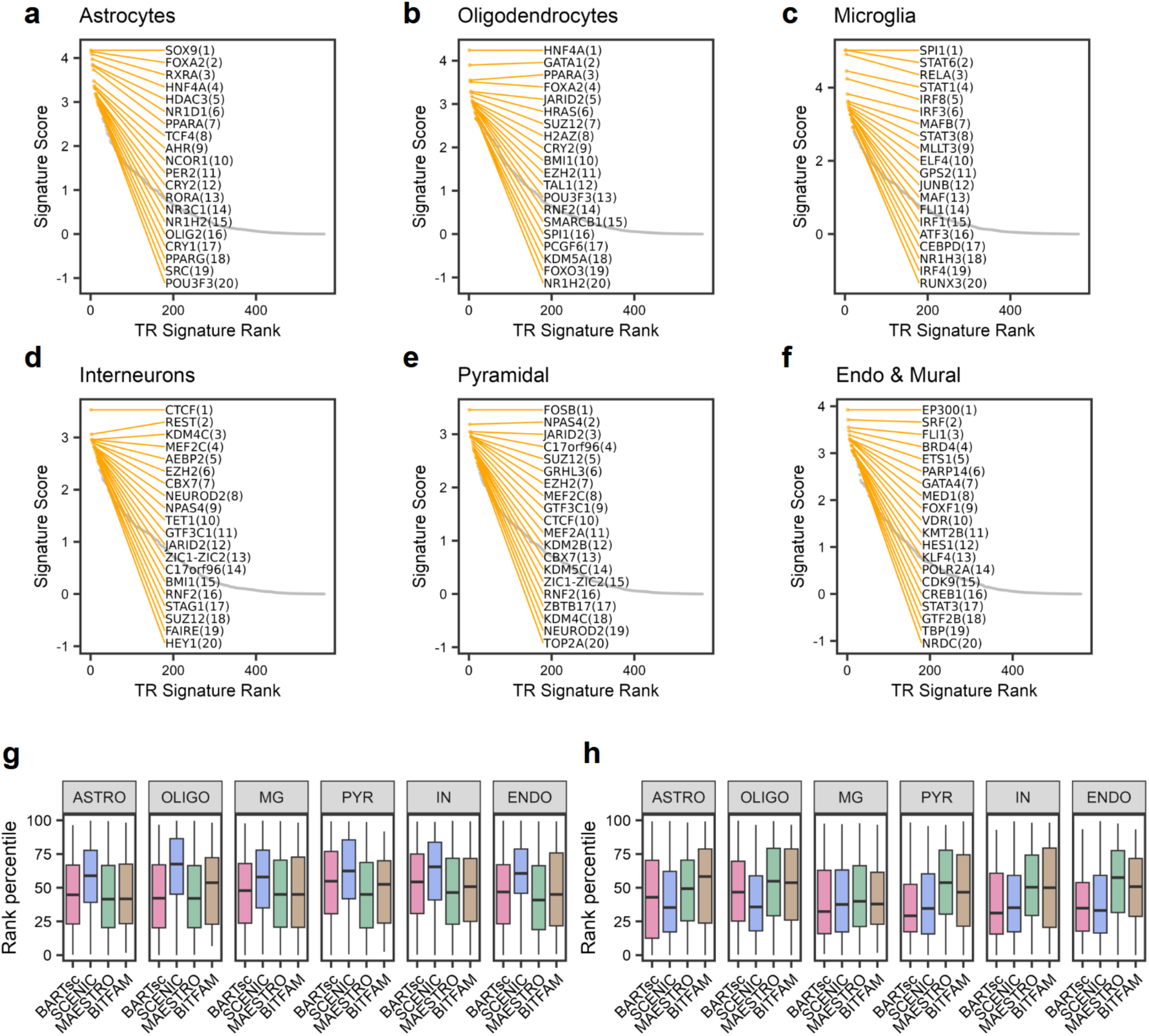
Signature score-based TR ranking and assessment of less-supported predictions across mouse cortex cell types. (**a**-**f**) Scatter plots showing TRs ranked by signature score across mouse cortex cell types. Top 20 TRs are highlighted and labeled with their rank in the parenthesis. (**g**, **h**) Distributions of the rank percentile of function-unknown TRs (**g**) and non-expressed TRs (**h**) predicted by each method across different cell types. In each box plot, the center line represents median; the box denotes the interquartile range (IQR); the whiskers extend to the minimum and maximum values within 1.5× IQR from the median.

**Figure S2.**
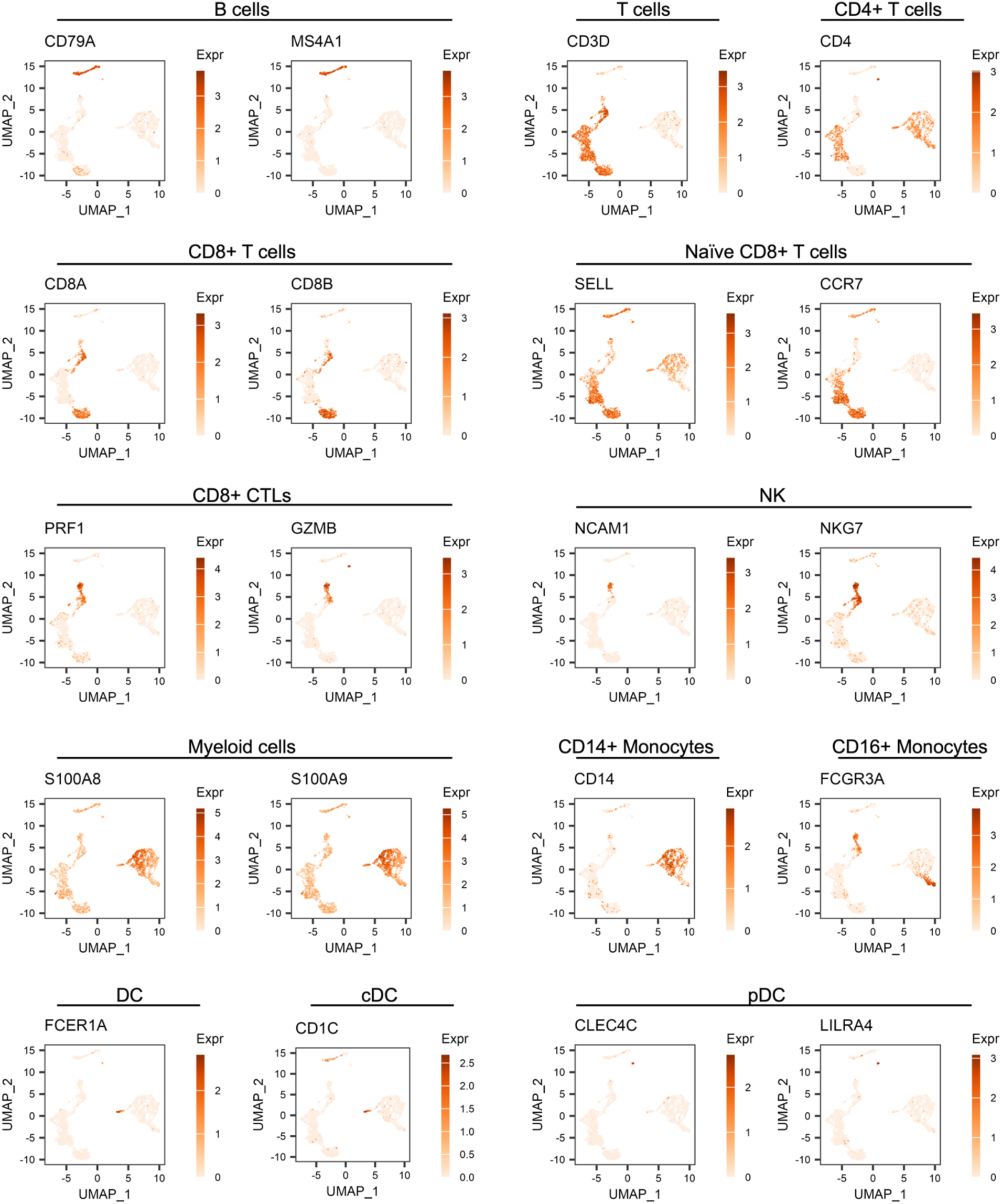
Marker gene expression across PBMC cell types. UMAP visualization of the PBMC scMultiome dataset, where each point represents a single cell colored by the normalized expression level of cell type marker genes.

**Figure S3.**
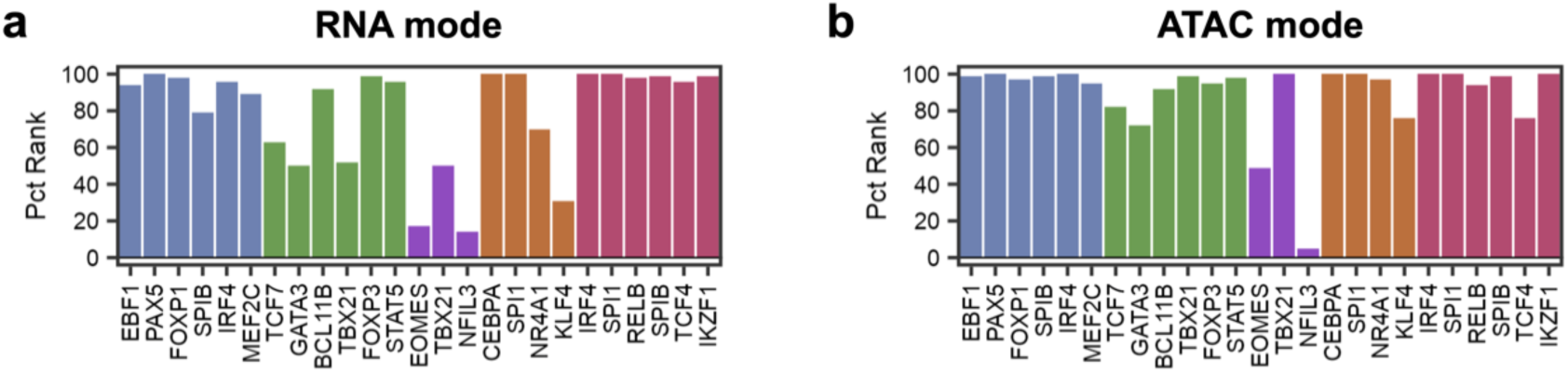
Unimodal signature scores of key PBMC regulators. (**a**, **b**) Bar plots showing percentile rank (PR) of known key regulators in the RNA mode (**a**), and ATAC mode (**b**). Bars are colored by the cell type in which the regulator is functionally associated.

**Figure S4.**
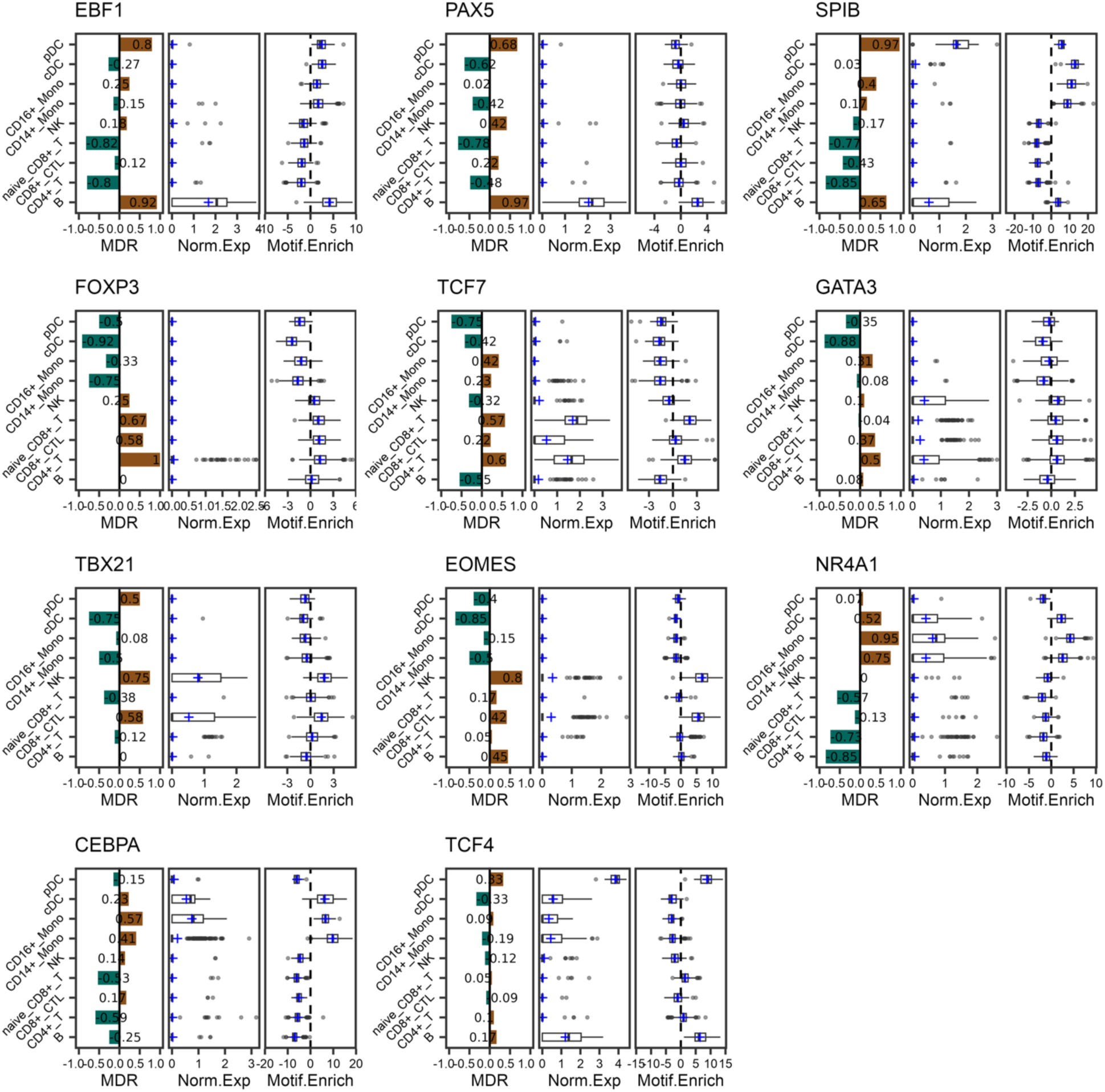
Relative TR activity is consistent with established cell-type-specific functions. Relative TR activity metrics for representative key regulators across PBMC cell types (B cells: EBF1, PAX5, SPIB; T cells: FOXP3, TCF7, GATA3, TBX21; NK cells: TBX21, EOMES; monocytes: NR4A1, CEBPA; dendritic cells: SPIB, TCF4). In each panel, Left: MDRs of the TR across PBMC cell types. Middle: distributions of normalized expression level of the TR across PBMC cell types. Right: distributions of the TR motif enrichment scores (chromVAR Z-score) across PBMC cell types. In the box plots, center lines represent medians; boxes denote the interquartile range (IQR); whiskers extending to the minimum and maximum values within 1.5× IQR from the median; and blue crosses representing mean values.

**Figure S5.**
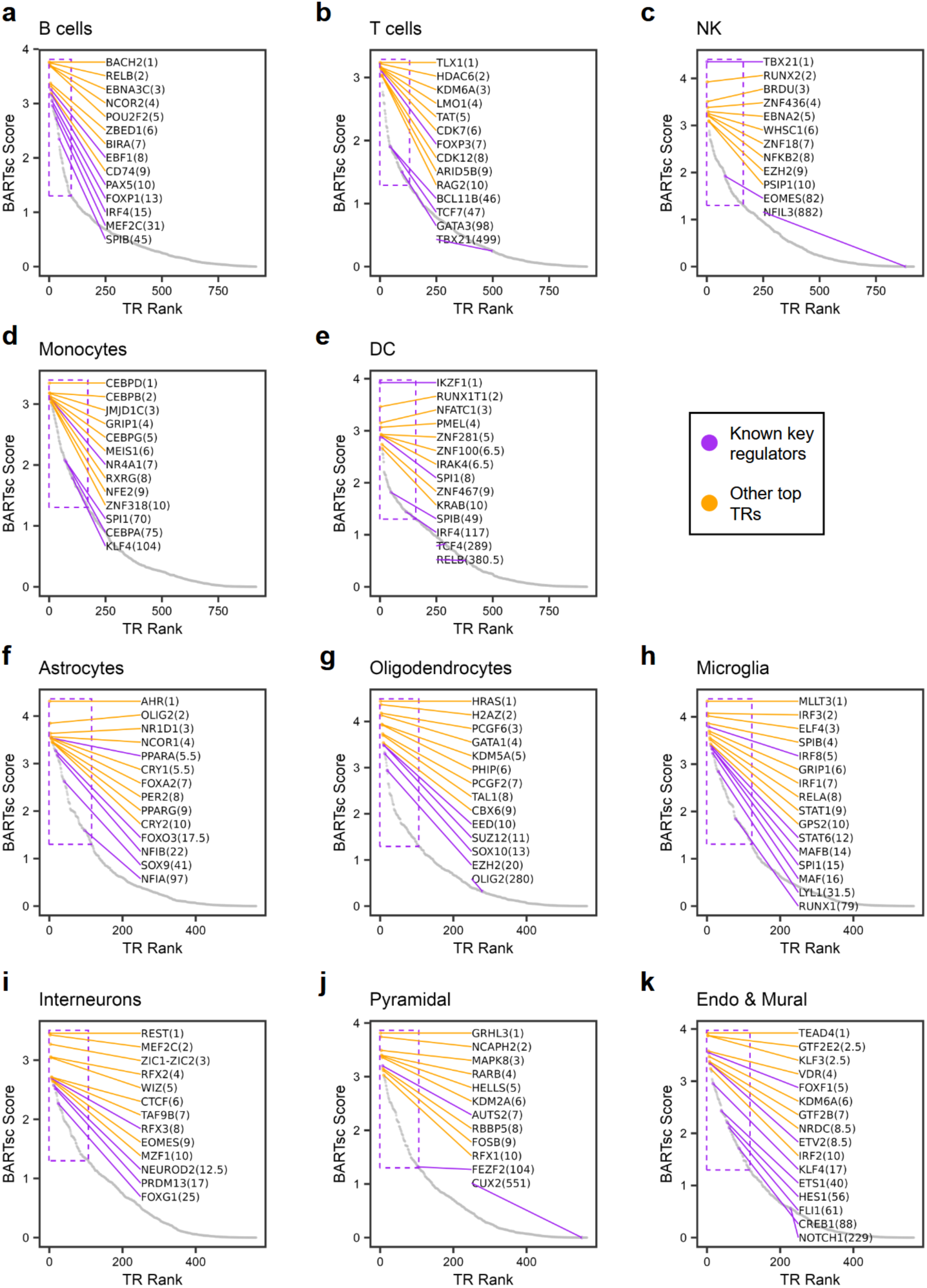
BARTsc identifies known and candidate key regulators across PBMC and cortex cell types. (**a**-**k**) Scatter plots showing TRs ranked by BARTsc score for human PBMC cell types (**a**-**e**) and mouse cortex cell types (**f**-**k**). The purple box indicates BARTsc-predicted key regulators with BARTsc scores above the default cutoff. Known key regulators, along with the other top 10 TRs are highlighted and labeled with their rank in the parenthesis.

**Figure S6.**
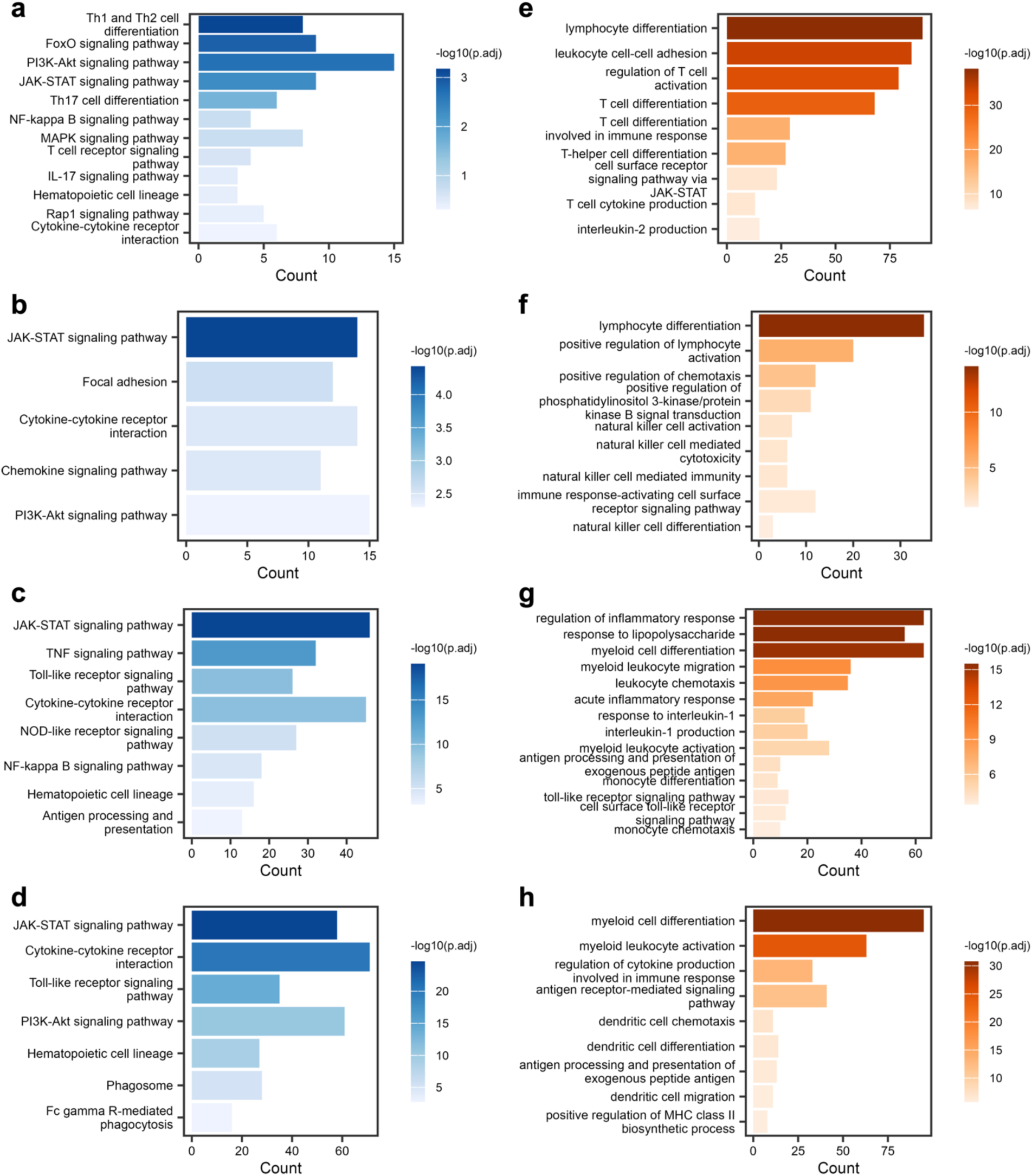
Functional enrichment of BARTsc-predicted key regulators and putative target genes across PBMC cell types. (**a**-**d**) Selected KEGG pathways enriched for BARTsc-predicted key regulators and their target genes across T cells (**a**), NK cells (**b**), monocytes (**c**), dendritic cells (**d**). (**e**-**h**) Selected Gene Ontology terms enriched for BARTsc-predicted key regulators and target genes across T cells (**e**), NK cells (**f**), monocytes (**g**), dendritic cells (**h**).

**Figure S7.**
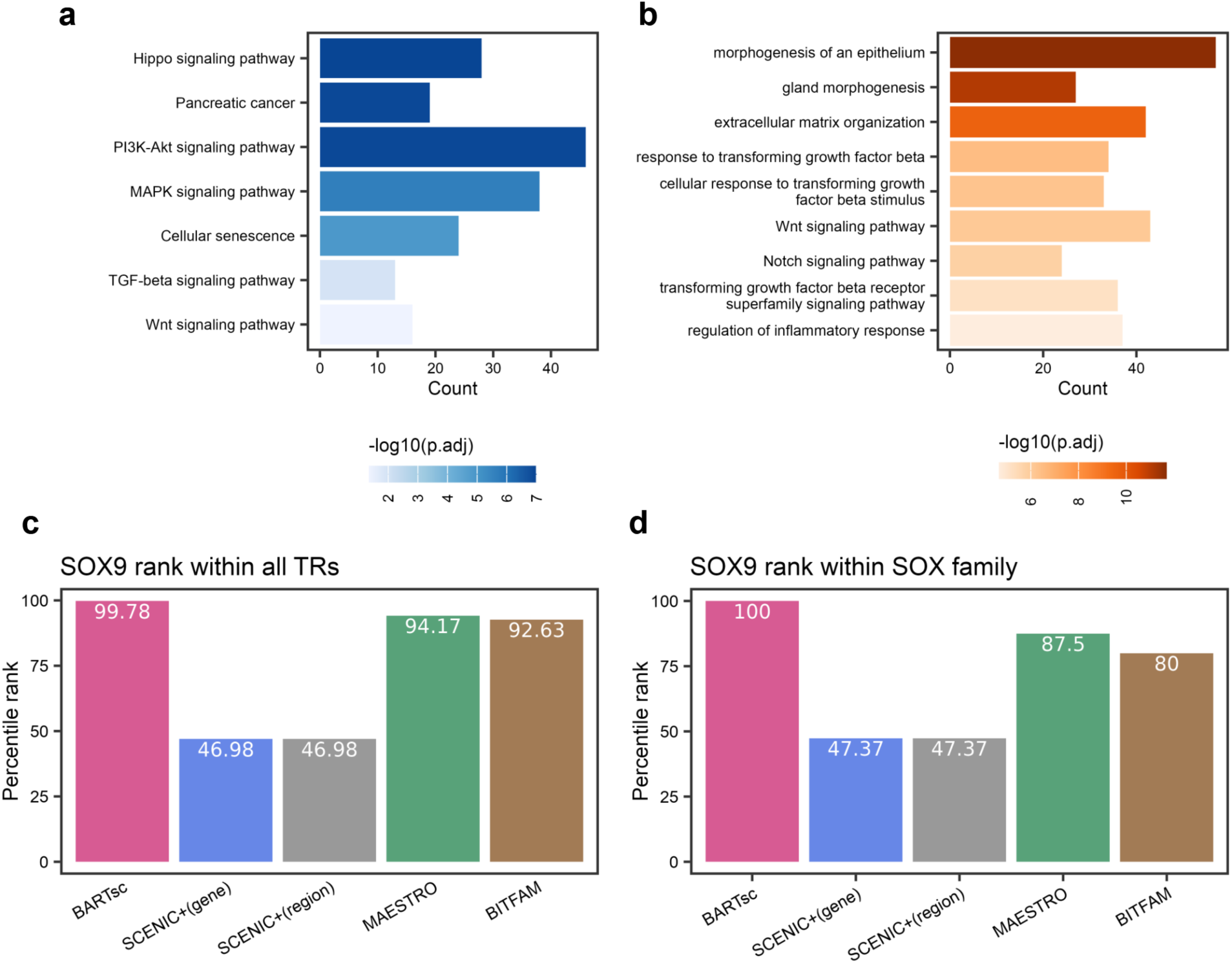
Functional characterization of ADM cells and assessment of SOX9 prediction. (**a**, **b**) Selected KEGG pathways (**a**) and Gene Ontology terms (**b**) enriched for BARTsc-predicted ADM cell key regulators and their target genes. (**c**, **d**) Percentile rank of SOX9 in the ranked list of TRs of ADM cells predicted by different methods. SOX9’s percentile rank was calculated among all TRs (**c**) and within the SOX family (**d**), respectively.

**Figure S8.**
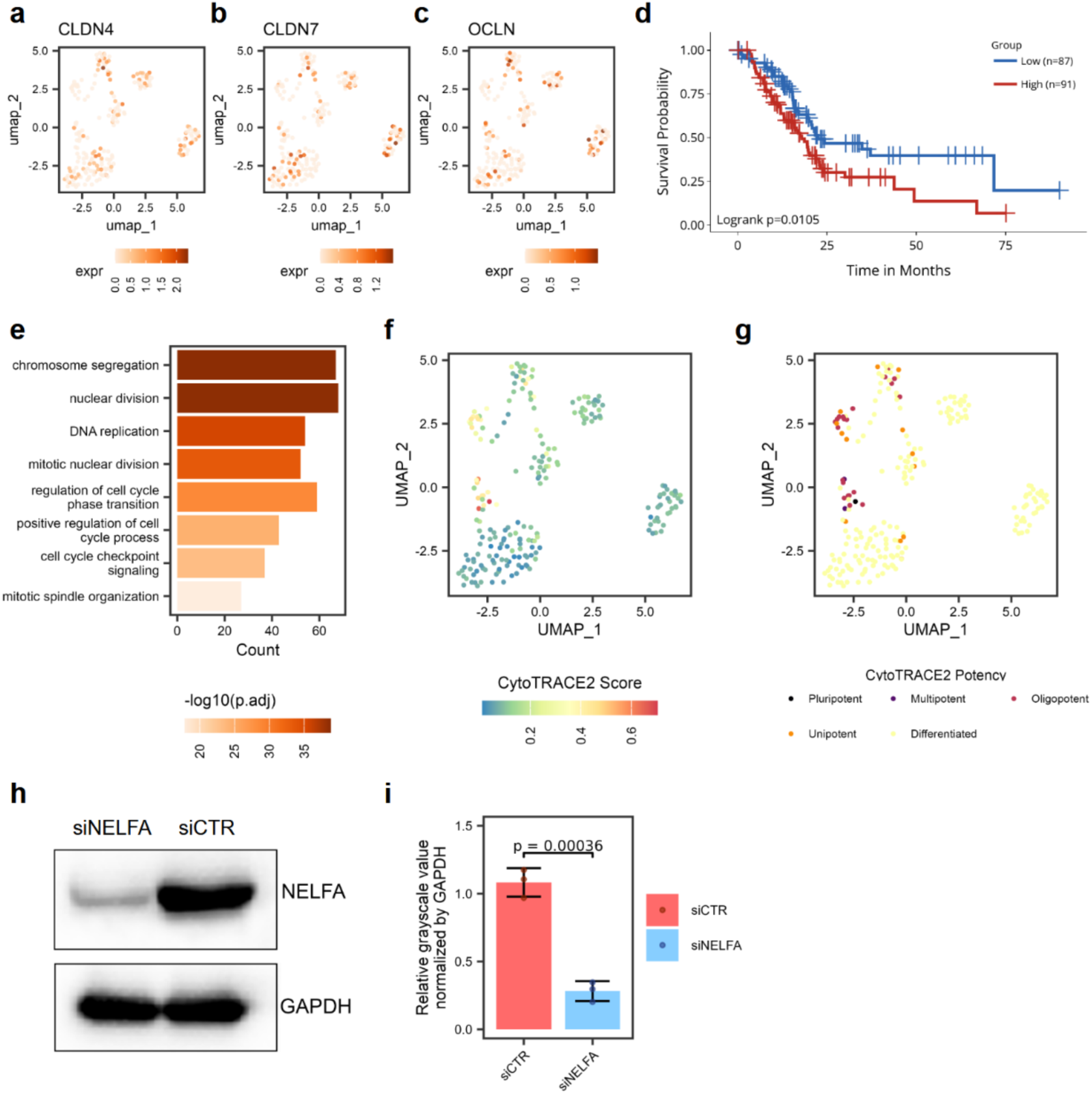
Regulatory and functional characterization of the c5 ductal cells in PDAC. (**a**-**c**) UMAP visualization of the PDAC scMultiome dataset, where each point represents a single cell colored by the normalized expression level of genes associated with tight junction: CLDN4 (**a**), CLDN7 (**b**) and OCLN (**c**). (**d**) Kaplan–Meier curves comparing PDAC patients with high versus low cluster5 (c5) signature expression (dataset used: TCGA-PAAD). Patients were stratified using the mean gene set enrichment score as the cutoff. P value was calculated by log-rank test. (**e**) Selected and Gene Ontology terms enriched for BARTsc-predicted c5 ductal cells key regulators and their target genes. (**f**, **g**) UMAP visualization of the PDAC scMultiome dataset, where each point represents a single cell colored by CytoTRACE2 score (**f**) and CytoTRACE2-predicted potency category (**g**). Higher CytoTRACE2 score indicates a less differentiated state. (**h, i**) Western blot analysis of NELFA protein expression in siNELFA and siControl groups, with GAPDH used as the loading control (**h**). Relative grayscale intensities were quantified and normalized to the loading control (**i**). Error bars indicate the standard deviation. P-value is calculated by one-sided *t*-test.

**Figure S9.**
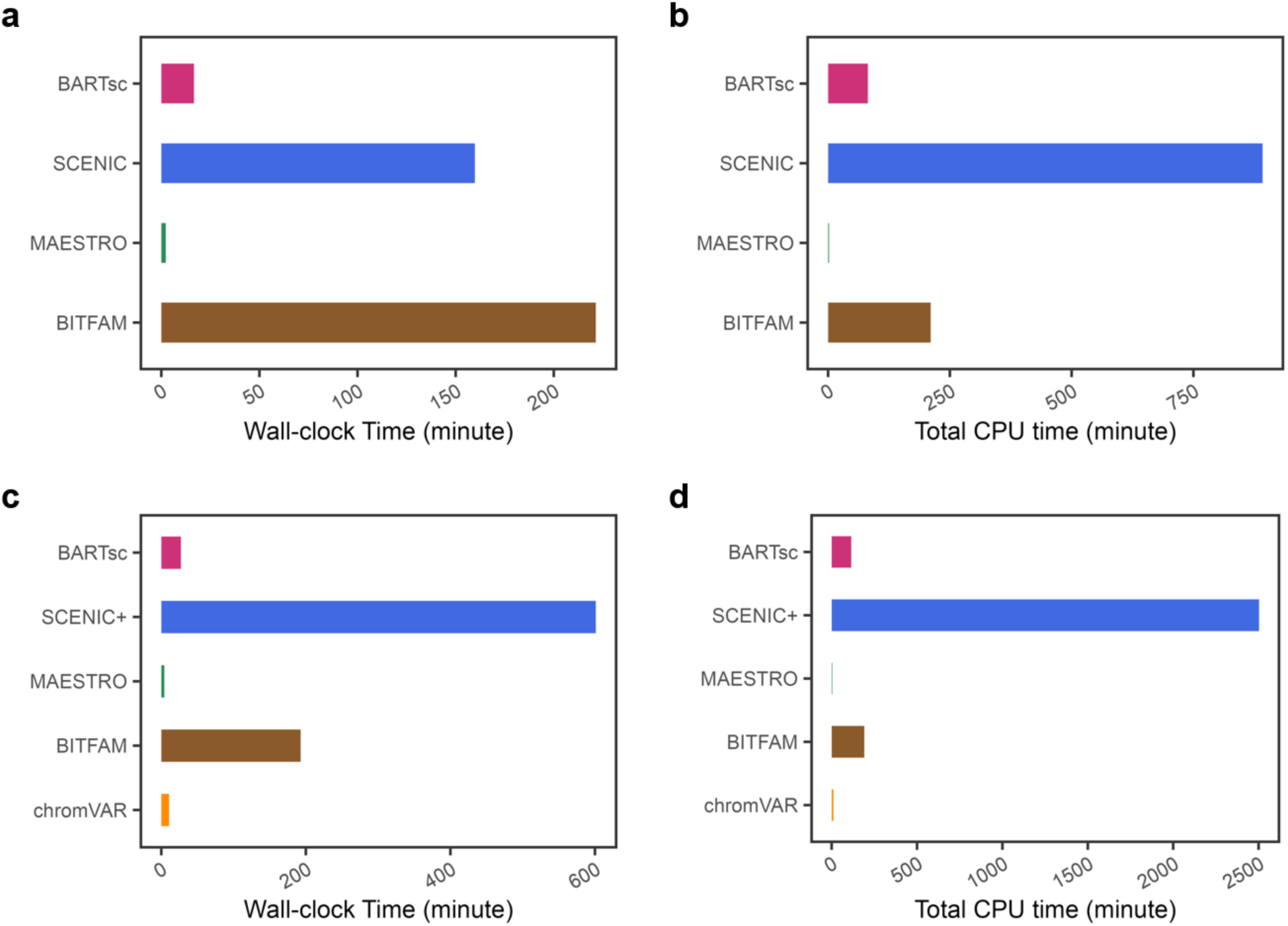
Runtime comparison of BARTsc and other methods. (**a**, **b**) Wall-clock time (**a**) and total CPU time (**b**) measured for the methods applied to the mouse cortex scRNA-seq dataset. (**c**, **d**) Wall-clock time (**c**) and total CPU time (**d**) used for the methods applied to analyze the human PBMC scMultiome dataset.

